# An Essential Role for *Argonaute 2* in EGFR-KRAS Signaling in Pancreatic Cancer Development

**DOI:** 10.1101/227264

**Authors:** Sunita Shankar, Jean Ching-Yi Tien, Ronald F. Siebenaler, Seema Chugh, Vijaya L. Dommeti, Sylvia Zelenka-Wang, Jessica Waninger, Kristin M. Juckette, Alice Xu, Xiao-Ming Wang, Malay Mody, Sanjana Eyunni, Andrew Goodrum, Grace Tsaloff, Yuping Zhang, Ingrid J. Apel, Lisha Wang, Javed Siddiqui, Richard D. Smith, Heather A. Carlson, John J. Tesmer, Xuhong Cao, Jiaqi Shi, Chandan Kumar-Sinha, Howard C. Crawford, Arul M. Chinnaiyan

**Author notes:** These authors contributed equally to this work (Sunita Shankar, Jean Ching-Yi Tien, Ronald F. Siebenaler, Seema Chugh). **Corresponding Author:** Arul M. Chinnaiyan, M.D., Ph.D., Investigator, Howard Hughes Medical Institute, American Cancer Society Professor, S. P. Hicks Endowed Professor of Pathology, Rogel Cancer Center, University of Michigan Medical School, 1400 E. Medical Center Dr. 5316 CCGC, Ann Arbor, MI 48109-0602.

## Abstract

KRAS and EGFR are known essential mediators of pancreatic cancer development. In addition, KRAS and EGFR have both been shown to interact with and perturb the function of Argonaute 2 (AGO2), a key regulator of RNA-mediated gene silencing. Here, we employed a genetically engineered mouse model of pancreatic cancer to define the effects of conditional loss of *AGO2* in *KRAS*^*G12D*^-driven pancreatic cancer. Genetic ablation of *AGO2* does not interfere with development of the normal pancreas or *KRAS*^*G12D*^-driven early precursor pancreatic intraepithelial neoplasia (PanIN) lesions. Remarkably, however, *AGO2* is required for progression from early to late PanIN lesions, development of pancreatic ductal adenocarcinoma (PDAC), and metastasis. *AGO2* ablation permits PanIN initiation driven by the EGFR-RAS axis, but rather than progressing to PDAC, these lesions undergo profound oncogene-induced senescence (OIS). Loss of *Trp53* (p53) in this model obviates the requirement of *AGO2* for PDAC development. In mouse and human pancreatic tissues, increased expression of AGO2 and elevated co-localization with RAS at the plasma membrane is associated with PDAC progression. Furthermore, phosphorylation of AGO2^Y393^ by EGFR disrupts the interaction of wild-type RAS with AGO2 at the membrane, but does not affect the interaction of mutant KRAS with AGO2. ARS-1620, a G12C-specific inhibitor, disrupts the KRAS^G12C^-AGO2 interaction specifically in pancreatic cancer cells harboring this mutant, demonstrating that the oncogenic KRAS-AGO2 interaction can be pharmacologically targeted. Taken together, our study supports a biphasic model of pancreatic cancer development: an *AGO2*-independent early phase of PanIN formation reliant on EGFR-RAS signaling, and an *AGO2*-dependent phase wherein the mutant KRAS-AGO2 interaction is critical to prevent OIS in PanINs and allow progression to PDAC.

## INTRODUCTION

*KRAS* mutations drive over 90% of pancreatic cancer, a disease with a dismal overall 5-year survival rate of only 9%^1^. Like all RAS GTPases KRAS is a molecular switch that transduces extracellular mitogenic signals by cycling between an active GTP-bound and an inactive GDP-bound state. Proteins that regulate the nucleotide loading of RAS, like GTPase activating proteins (GAPs) or guanine exchange factors (GEFs) recruit RAS to the plasma membrane in response to activated (i.e., phosphorylated) growth factor receptors, like EGFR^2-4^. Recurrent oncogenic driver mutations in *RAS* decrease terminal phosphate cleavage, resulting in the accumulation of its active GTP-bound form at the plasma membrane. Constitutively active RAS leads to aberrant signaling activities through interactions with multiple effector proteins at the plasma membrane^2,3,5^.

Genetically engineered mouse models (GEMMs) of pancreatic cancer were developed by expression of a single oncogenic *KRAS*^*G12D*^ allele in the mouse exocrine pancreas. In this model, pre-invasive pancreatic intraepithelial (PanINs) lesions progress to pancreatic adenocarcinoma (PDAC) reflective of the human disease^6^. Further, the pathophysiology and site of metastases observed in this model closely mimic human pancreatic cancer, underscoring its applicability for studying the human disease. Use of such GEMMs have been instrumental in defining the key events that characterize PanIN development and PDAC progression^7-9^. Of particular relevance is the observation that EGFR, the upstream modulator of RAS signaling, is essential for *KRAS*^*G12D*^-driven PanIN development^10,11^, despite mutual exclusivity of *EGFR* and *RAS* aberrations in cancer. However, the requirement for EGFR at the early stage of PanIN development has not translated to successful treatment of pancreatic cancer^12,13^. Despite our understanding of the signaling events triggered by oncogenic RAS, targeting KRAS remains a challenging prospect^4,14^.

To investigate potential additional modulators of RAS-mediated oncogenesis, we previously performed a screen for direct interactors of RAS in a panel of cancer cell lines and identified a direct interaction between KRAS and Argonaute 2 (AGO2), independent of *KRAS* mutation status^15^. The N-terminal domain of AGO2 was found to bind the regulatory Switch II region in RAS and was required for oncogenic *KRAS*-driven cellular transformation. Notably, AGO2 RNA silencing activity was inhibited in cells expressing mutant RAS compared to wild-type RAS, suggesting that KRAS binding, and not the RNAi function of AGO2, was essential for oncogenesis. Interestingly, Shen et al. had previously identified a functional interaction between EGFR and AGO2^16^. Their study showed that EGFR phosphorylates AGO2 at tyrosine 393 under hypoxic stress to alter its RNAi function.

Given these previous observations, we have employed established mouse models of pancreatic cancer to determine the *in vivo* requirement of *AGO2* in pancreatic cancer development in the context of KRAS signaling. Our data show that oncogenic *KRAS*-initiated PanIN formation is reliant on EGFR and wild-type RAS signaling, independent of *AGO2*. Strikingly, however, we identify a critical dependence on *AGO2* for PanIN progression to PDAC, bypassed by loss of *p53*. While defining an essential role for *AGO2* in PDAC progression, we also further our understanding of how the KRAS-AGO2 interaction is regulated through EGFR activation. Disruption of the oncogenic KRAS-AGO2 association may, therefore, represent a novel point of therapeutic intervention to prevent pancreatic cancer progression.

## RESULTS

### Loss of *AGO2* does not affect normal pancreas development or early stage PanIN formation in *KRAS*^*G12D*^ mice

To investigate the role of *AGO2* in the development of pancreatic cancer *in vivo*, we employed the genetically engineered mouse model of pancreatic cancer initiated by a conditionally activated allele of *KRAS* (ref.^6^), *KRAS*^*LSL-G12D/+*^ (KRAS^G12D^, shown in **Fig. 1a**). Crossing *KRAS*^*G12D*^ mice with animals harboring *Cre* recombinase knocked into the pancreas-specific promoter, *p48* (*p48Cre*), yields *KRAS*^*G12D*^;*p48Cre* mice that develop pancreatic intraepithelial neoplasia (PanINs) precursor lesions beginning around 8 weeks of age^6^. Over time, these PanINs progress to pancreatic ductal adenocarcinoma (PDAC) and develop metastases, faithfully mimicking the human disease. In order to evaluate the potential consequences of *AGO2* ablation in this model, we generated transgenic mice with both *KRAS*^*G12D*^ and conditionally deleted allele(s) of *AGO2* (ref.^17^) (**Fig. 1a**). The resulting *KRAS*^*G12D*^;*p48Cre* mice were either wild-type, heterozygous, or homozygous for the conditional allele of *AGO2* (hereafter referred to as *AGO2*^*+/+*^;*KRAS*^*G12D*^;*p48Cre, AGO2*^*fl/+*^;*KRAS*^*G12D*^;*p48Cre,* and *AGO2*^*fl/fl*^;*KRAS*^*G12D*^;*p48Cre,* respectively). Genomic PCR confirmed Cre-driven excision and recombination of the oncogenic *KRAS* allele^6^ in pancreata from mice with *KRAS*^*G12D*^; *p48Cre* alleles (**Supplementary Fig. 1a**). Further, qRT-PCR analysis showed significant reduction in *AGO2* expression in *AGO2*^*fl/fl*^;*KRAS*^*G12D*^;*p48Cre* mice (**Supplementary Fig. 1b**), confirming Cre-mediated mutant *KRAS* activation with concomitant loss of *AGO2* expression in the pancreas.

**Figure 1.**
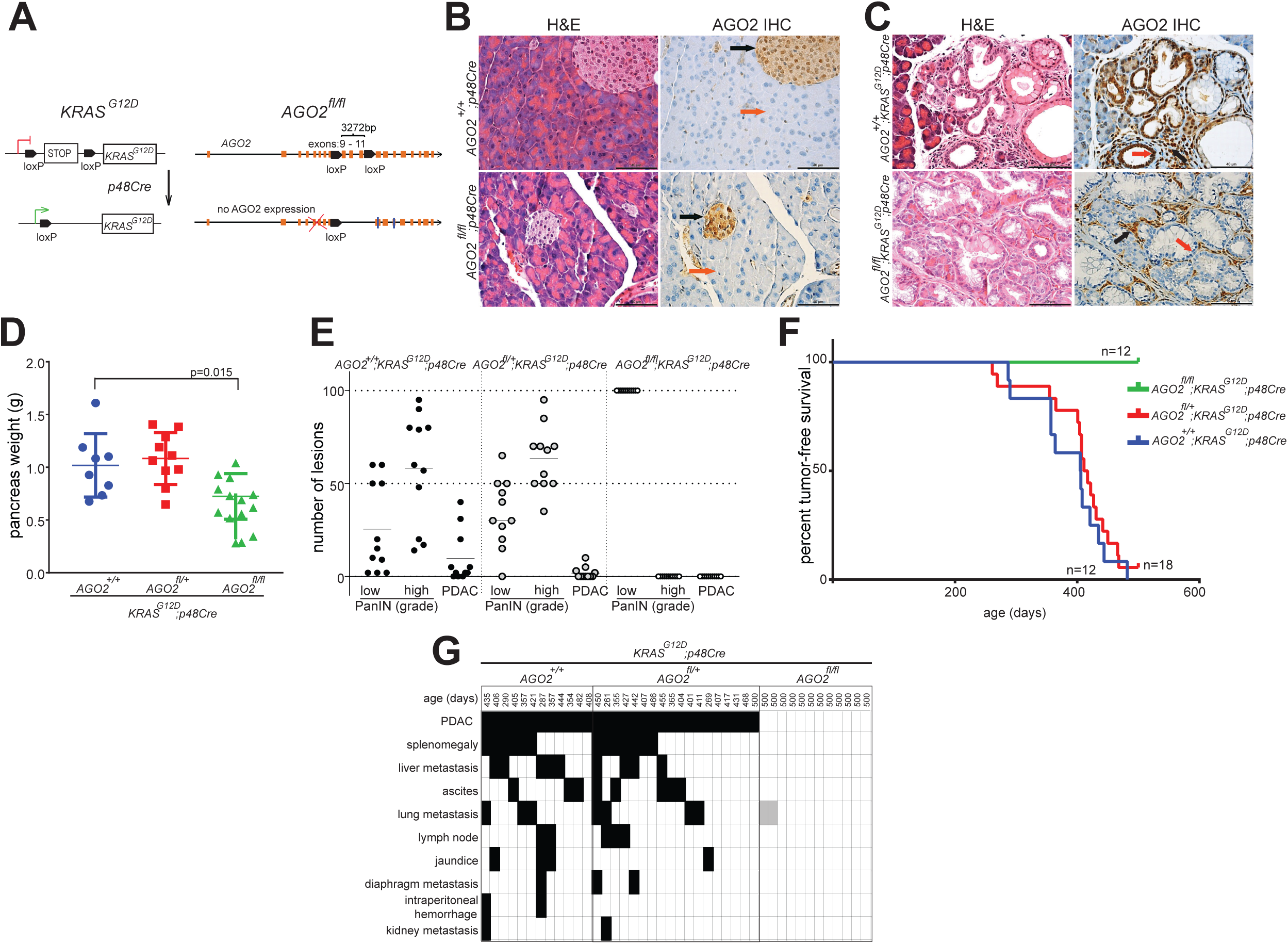
*AGO2* is essential for progression of precursor PanIN lesions to PDAC. (A) Schematic of the conditionally activated endogenous alleles of *KRAS*^*G12D*^ and *AGO2* used in the study to generate the *AGO2*^*fl/fl*^;*KRAS*^*G12D*^;*p48Cre* experimental mice. (B) Representative images of H&E and AGO2 IHC analysis of pancreata obtained from *AGO2*^*+/+*^;*p48Cre* and *AGO2*^*fl/fl*^;*p48Cre* genotypes. Orange and black arrows indicate AGO2 expression in acinar cells and islets of Langerhans, respectively. (C) Representative H&E and IHC analysis for AGO2 in pancreata obtained from 12-week old mice from the *AGO2*^*+/+*^;*KRAS*^*G12D*^;*p48Cre* and *AGO2*^*fl/fl*^;*KRAS*^*G12D*^;*p48Cre* genotypes. Orange and black arrows indicate AGO2 staining in the PanIN and stromal regions, respectively. (D) Scatter plot showing the weight of pancreata obtained from mice of the indicated genotypes aged over 400 days. (E) Histogram showing average number of early and late PanIN lesions observed in 11 *AGO2*^*+/+*^;*KRAS*^*G12D*^;*p48Cre*, 11 *AGO2*^*fl/+*^;*KRAS*^*G12D*^;*p48Cre*, and 11 *AGO2*^*fl/fl*^;*KRAS*^*G12D*^;*p48Cre* mice aged over 400 days. For *AGO2*^*fl/fl*^;*KRAS*^*G12D*^;*p48Cre* mice, only lesions that do not express AGO2 have been included. (F) Kaplan-Meier curve for tumor-free survival of *AGO2*^*+/+*^;*KRAS*^*G12D*^;*p48Cre, AGO2*^*fl/+*^;*KRAS*^*G12D*^;*p48Cre*, and *AGO2*^*fl/fl*^; *KRAS*^*G12D*^;*p48Cre* mice aged over 500 days. (G) Chart showing PDAC (within the pancreas), the different metastatic lesions, and abnormal pathologies (black boxes) observed in each mouse of the indicated genotypes aged over 500 days. Gray boxes in the *AGO2*^*fl/fl*^;*KRAS*^*G12D*^;*p48Cre* group indicate abnormal pathology observed at the indicated site and are addressed in further detail in **Supplementary Fig. 5**.

Histological analysis of pancreata from mice with Cre-mediated *AGO2* ablation (*AGO2*^*fl/fl*^; *p48Cre*) showed normal morphology (**Fig. 1b**, left panels) with no differences in pancreatic weight compared to pancreata from *AGO2*^*+/+*^;*p48Cre* mice (**Supplementary Fig. 1c**). This suggests that loss of *AGO2* in the acinar cells of the exocrine compartment does not grossly interfere with pancreas development. Immunohistochemical (IHC) staining with a monoclonal antibody specific to AGO2 (**Supplementary Fig. 2** and **Supplementary Table 1**) showed minimal expression of AGO2 in the acinar cells of both *AGO2*^*+/+*^;*p48Cre* and *AGO2*^*fl/fl*^;*p48Cre* pancreata (**Fig. 1b**, right panels). Interestingly, relatively higher expression of AGO2 was seen in pancreatic endocrine cells (islets of Langerhans), which was unaffected by the acinar cell-specific ablation of *AGO2*. These data indicate a non-essential role for *AGO2* in the acinar cells during normal pancreatic development. However, expression of *KRAS*^*G12D*^ in the pancreatic acinar cells led to increased AGO2 expression in the PanINs as well as the surrounding stroma in 12-week old *AGO2*^*+/+*^;*KRAS*^*G12D*^;*p48Cre* mice (**Fig. 1c**, top panels). Notably, we observed PanIN lesions in *AGO2*^*fl/fl*^;*KRAS*^*G12D*^;*p48Cre* pancreata lacking *AGO2* expression (**Fig. 1c**, lower panels). These early stage precursor PanIN lesions in *AGO2*^*fl/fl*^;*KRAS*^*G12D*^;*p48Cre* pancreata were morphologically indistinguishable from those arising in *AGO2*^*+/+*^;*KRAS*^*G12D*^;*p48Cre* mice. While PanIN lesions in *AGO2*^*fl/fl*^;*KRAS*^*G12D*^;*p48Cre* mice do not exhibit AGO2 expression, the surrounding stromal cells continue to express relatively high levels of AGO2. Further, PanINs from both *AGO2*^*+/+*^;*KRAS*^*G12D*^;*p48Cre* and *AGO2*^*fl/fl*^;*KRAS*^*G12D*^;*p48Cre* mice displayed high mucin content as seen by Alcian blue staining^18^ and similar gross weights of the pancreas, indicating indistinct phenotypes at the 12-week time point (**Supplementary Fig. 3a-b**).

**Figure 2.**
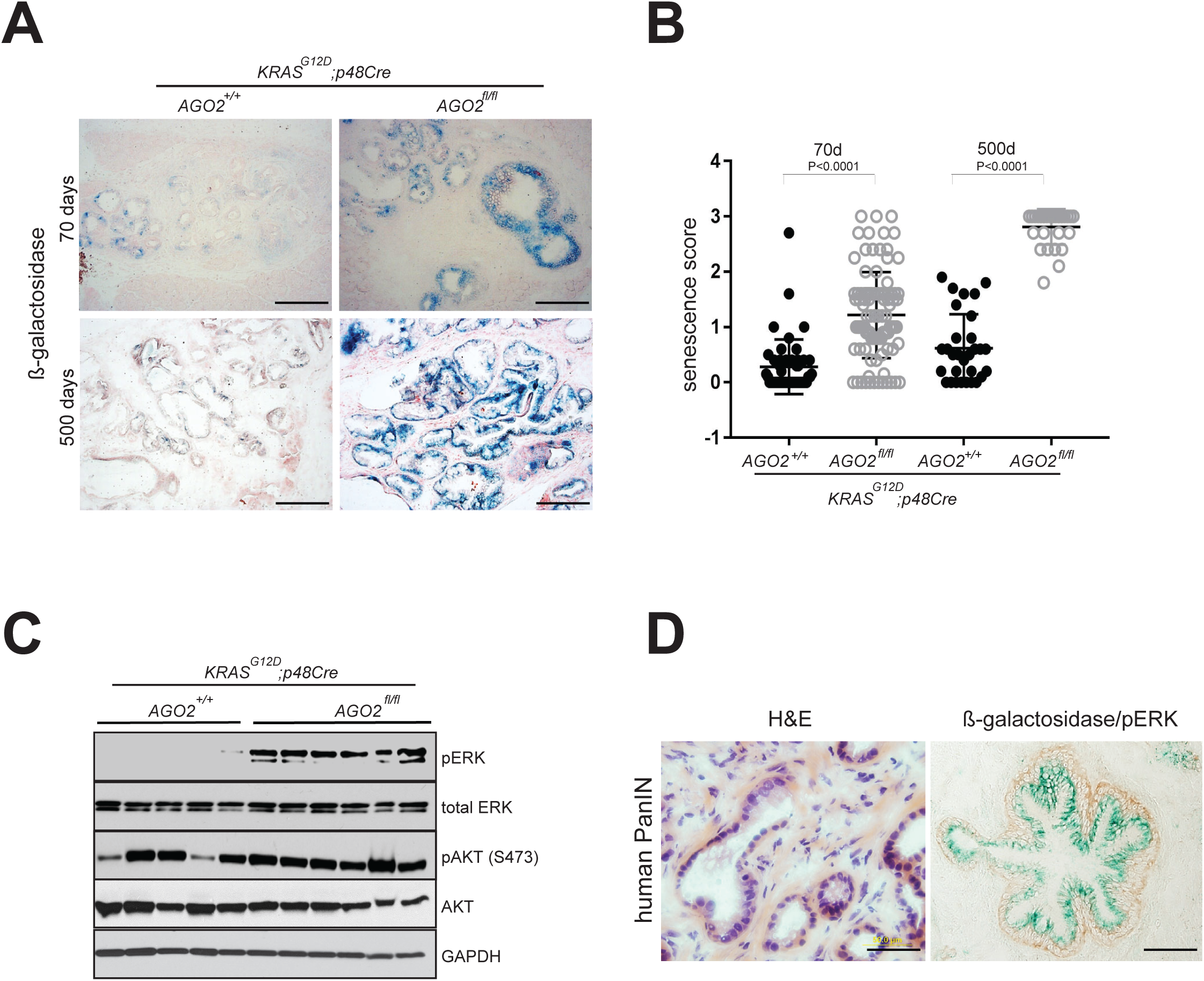
Oncogene-induced senescence upon *AGO2* loss prevents PanIN to PDAC progression. (A) β-galactosidase staining of pancreatic sections from *AGO2*^*+/+*^;*KRAS*^*G12D*^;*p48Cre* and *AGO2*^*fl/fl*^;*KRAS*^*G12D*^;*p48Cre* at 70- and 500-day time points. Scale bar, 100 µm. (B) Scatter plot showing β-galactosidase staining in low grade PanINs observed in at least three mice of the indicated genotypes at 70- or 500-day time points. Intensity of staining and percent cells within 30 low grade PanINs were used to determine the senescence score = intensity x percent positive cells. p values were determined using a t-test. (C) Immunoblot analysis of RAS-driven MAPK (indicated by pERK) and PI3K (indicated by pAKT) signaling from individual pancreata obtained from mice of the indicated genotypes, aged to 400 days. (D) Representative images of H&E staining (left) and dual staining for β-galactosidase and phospho-ERK (right) in human pancreatic tissue with PanINs (representative staining of at least 10 PanINs from two patients).

**Figure 3.**
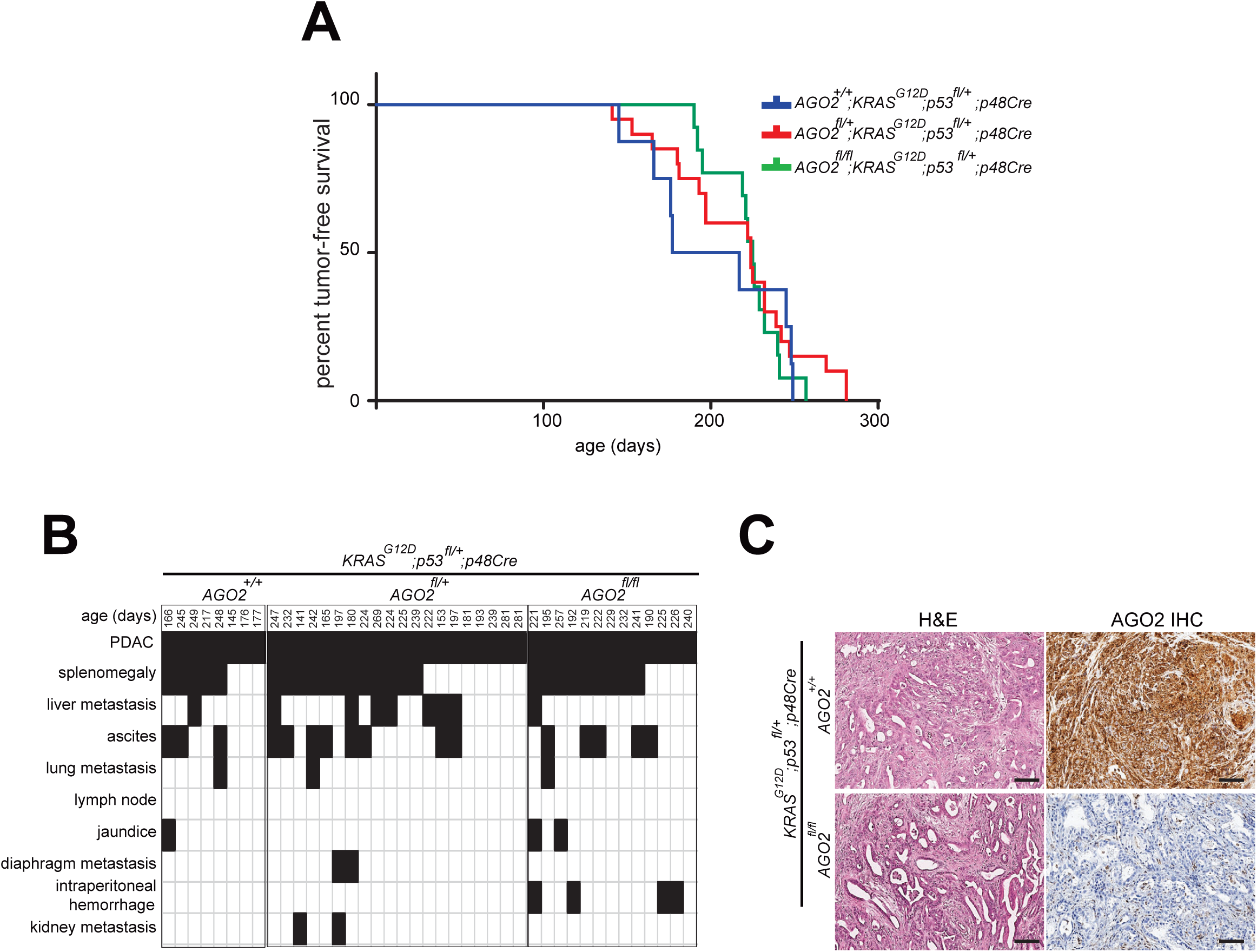
*AGO2* is not required for PDAC progression in the *Kras*^*G12D*^;*Trp53*^*fl/+*^;*p48Cre* (KPC) mouse model. (A) Kaplan-Meier tumor-free survival of Ago*2*^*+/+*^;*Kras*^*G12D*^;*Trp53*^*fl/+*^;*p48Cre,* Ago*2*^*fl/+*^;*Kras*^*G12D*^;*Trp53*^*fl/+*^;*p48Cre*, and *Ago2*^*fl/fl*^;*Kras*^*G12D*^;*Trp53*^*fl/+*^;*p48Cre* mice. (B) Chart showing PDAC (within the pancreas), the different metastatic lesions, and abnormal pathologies (black boxes) observed in each mouse of the indicated genotypes. (C) Representative H&E and AGO2 IHC in the indicated genotype.

### Loss of *AGO2* in *KRAS*^*G12D*^ mice blocks pancreatic cancer development, metastatic progression, and lethality

Surprisingly, over a longer course of time, mice aged over 400 days (57 weeks) showed significantly increased pancreatic weights in both the *AGO2*^*+/+*^;*KRAS*^*G12D*^;*p48Cre* and *AGO2*^*fl/+*^;*KRAS*^*G12D*^;*p48Cre* cohort compared to *AGO2*^*fl/fl*^;*KRAS*^*G12D*^;*p48Cre* mice, suggestive of a higher tumor burden in mice with at least one functional allele of *AGO2* (**Fig. 1d**). Histological analysis of pancreata at the 400-day time point showed early/late PanIN lesions and some PDAC development in *AGO2*^*+/+*^;*KRAS*^*G12D*^;*p48Cre* and *AGO2*^*fl/+*^;*KRAS*^*G12D*^;*p48Cre* mice with a distribution consistent with those previously reported^11,19^. However, in the *AGO2*^*fl/fl*^;*KRAS*^*G12D*^;*p48Cre* mice, mostly early stage PanIN lesions were observed, strikingly, with no evidence of PDAC (**Fig. 1e**). Occasionally, higher grade PanIN lesions were also observed in *AGO2*^*fl/fl*^;*KRAS*^*G12D*^;*p48Cre* pancreata, but these lesions invariably showed AGO2 expression (**Supplementary Fig. 4**), indicative of likely escape from Cre recombination, as has been previously noted in other contexts^11,20^.

**Figure 4.**
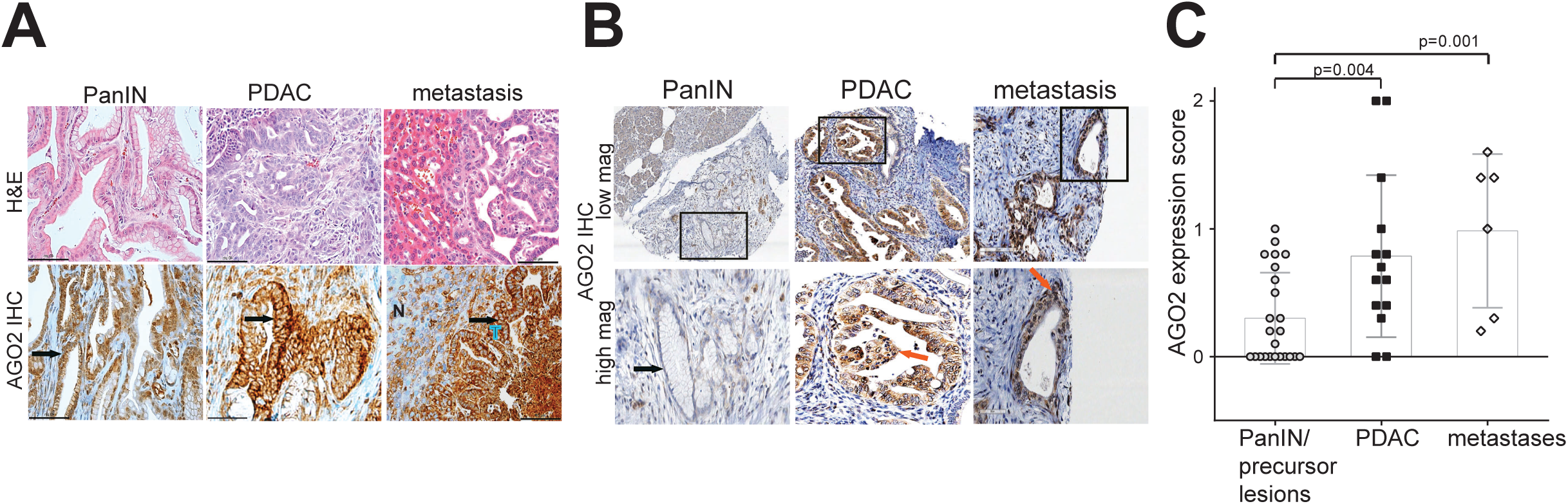
Increased AGO2 protein expression during mouse and human PDAC progression. (A) Representative images of AGO2 IHC analysis within an individual *AGO2*^*+/+*^;*KRAS*^*G12D*^;*p48Cre* mouse showing increased AGO2 expression in PDAC and metastasis compared to PanIN lesions. Arrows point to PanIN, PDAC, or metastatic PDAC in respective panels. In the metastasis panel, N=normal liver and T=tumor. Scale bar, 40 µm. (B) Representative images of IHC analysis for AGO2 expression in human PDAC progression showing elevated AGO2 protein expression in PDAC and metastatic tissue. Lower panels show higher magnifications of areas marked in the upper panels. Low and high magnification indicate 8X and 20X images, respectively. Arrows point to PanIN and PDAC. (C) Box and scatter plot showing AGO2 expression on a human tissue microarray (TMA) containing 24 precancerous, 14 PDAC, and 6 metastatic PDAC lesions, as determined by IHC analysis. Each sample was scored for intensity of stain and percent tumor cells staining for AGO2, and the final score = intensity x percent positive cells. p values were determined using a t-test.

In order to further examine the effect of *AGO2* loss on tumor-free survival, a cohort of transgenic mice was monitored over 500 days. Twelve of 12 *AGO2*^*+/+*^;*KRAS*^*G12D*^;*p48Cre* and 18 of 19 *AGO2*^*fl/+*^;*KRAS*^*G12D*^;*p48Cre* mice died over a median of 406 and 414 days, respectively, typical for a murine model expressing *KRAS*^*G12D*^ in the pancreas^7,9,21^. Remarkably, however, all 12 of 12 mice with homozygous *AGO2* deficiency (*AGO2*^*fl/fl*^;*KRAS*^*G12D*^;*p48Cre)* had survived at the cut-off time point of 500 days (**Fig. 1f**). PDAC was observed in pancreata of all mice that expressed *AGO2* (wild-type or heterozygous expression), but mice deficient for *AGO2* developed only early PanIN precursor lesions without progression to PDAC (**Fig. 1g**). Necropsies of experimental mice from the different genotypes were also assessed for grossly visible metastases and abnormal pathologies^22^ and revealed frequent metastases in the *AGO2*^*+/+*^;*KRAS*^*G12D*^;*p48Cre* and *AGO2*^*fl/+*^;*KRAS*^*G12D*^;*p48Cre* genotypes, but *AGO2*^*fl/fl*^;*KRAS*^*G12D*^;*p48Cre* mice rarely showed abnormal pathologies and never frank adenocarcinoma or metastases (**Fig. 1g**). While all metastatic lesions from mice expressing *AGO2* showed PDAC, analyses of lungs with abnormal pathologies in two of the *AGO2*^*fl/fl*^;*KRAS*^*G12D*^;*p48Cre* mice (marked as gray boxes) showed a single benign lesion each, associated with AGO2 expression (representing lesions with non-recombined alleles/non-pancreatic origin) without indication of PDAC (**Supplementary Table 2** and **Supplementary Fig. 5**). One mouse of the *AGO2*^*fl/fl*^;*KRAS*^*G12D*^;*p48Cre* genotype developed a pancreatic cyst (without AGO2 expression), histologically resembling the mucinous cystic neoplasm, and survived for 368 days (**Supplementary Table 2** and **Supplementary Fig. 5**). Taken together, these data show that *AGO2* is not essential for either normal pancreatic development or *KRAS*^*G12D*^-driven PanIN formation. Notably, however, *AGO2* is indispensable for progression of PanINs to PDAC, despite the presence of other Argonaute proteins not deleted in this model with compensatory and overlapping RNAi functions.

**Figure 5.**
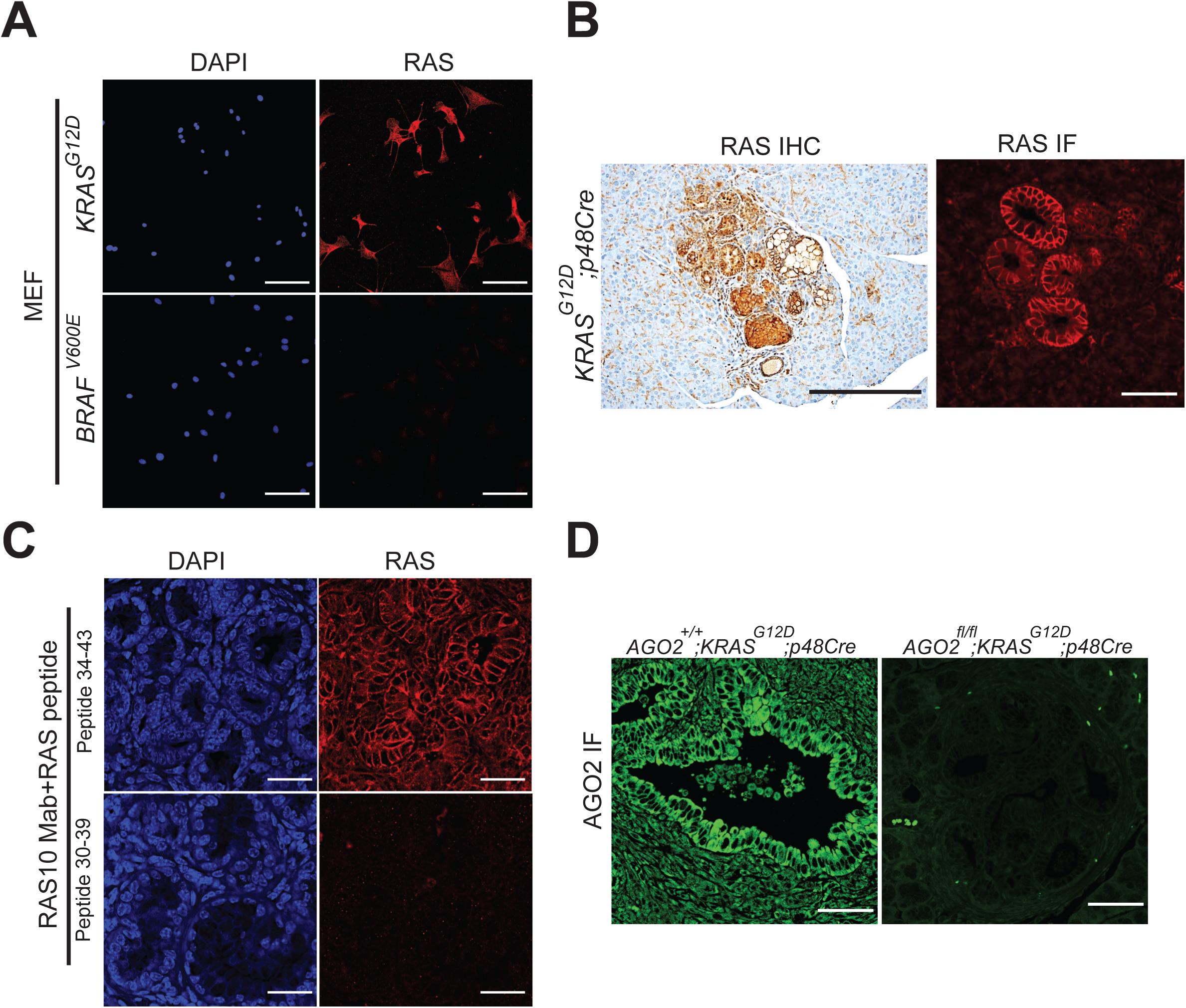
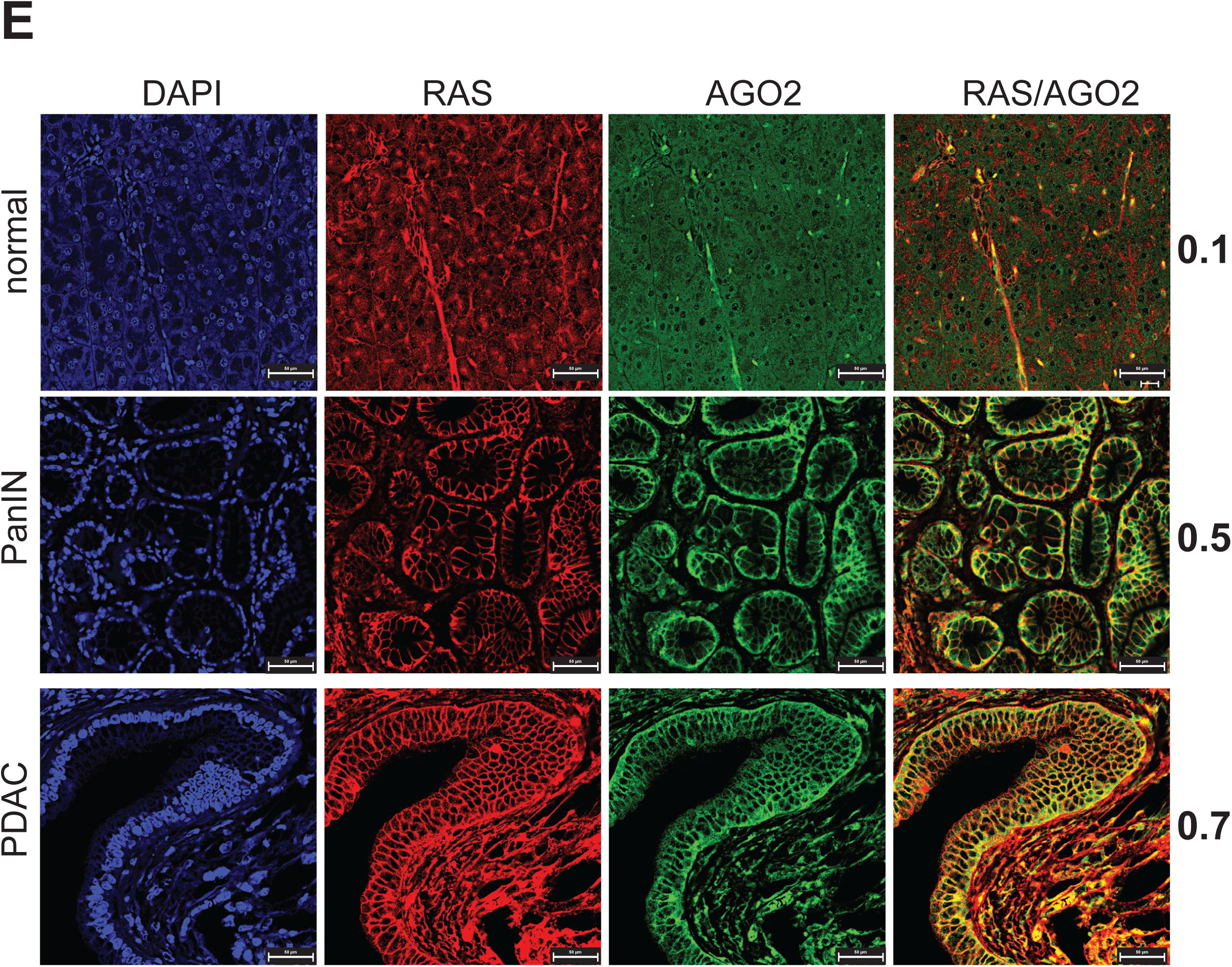
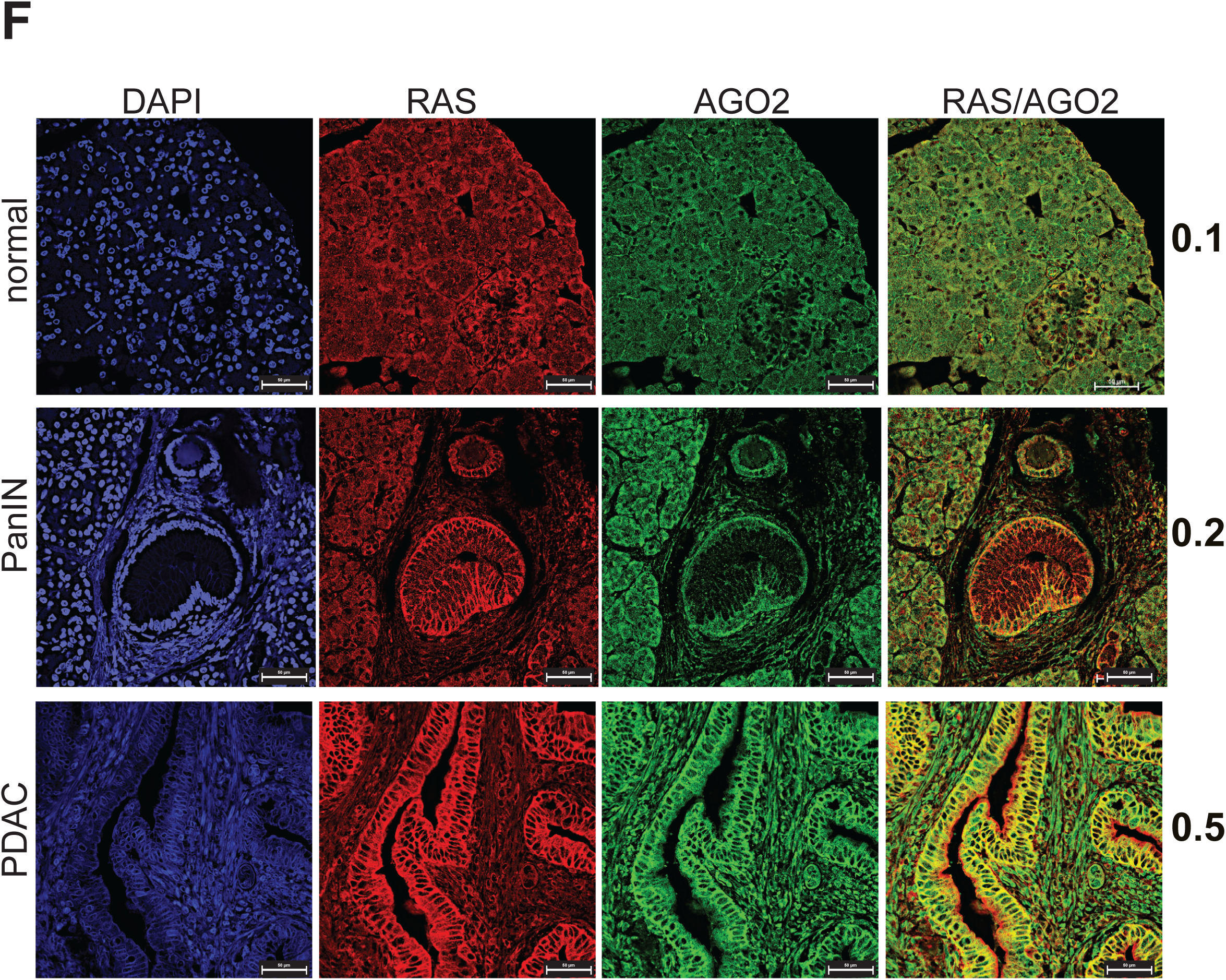
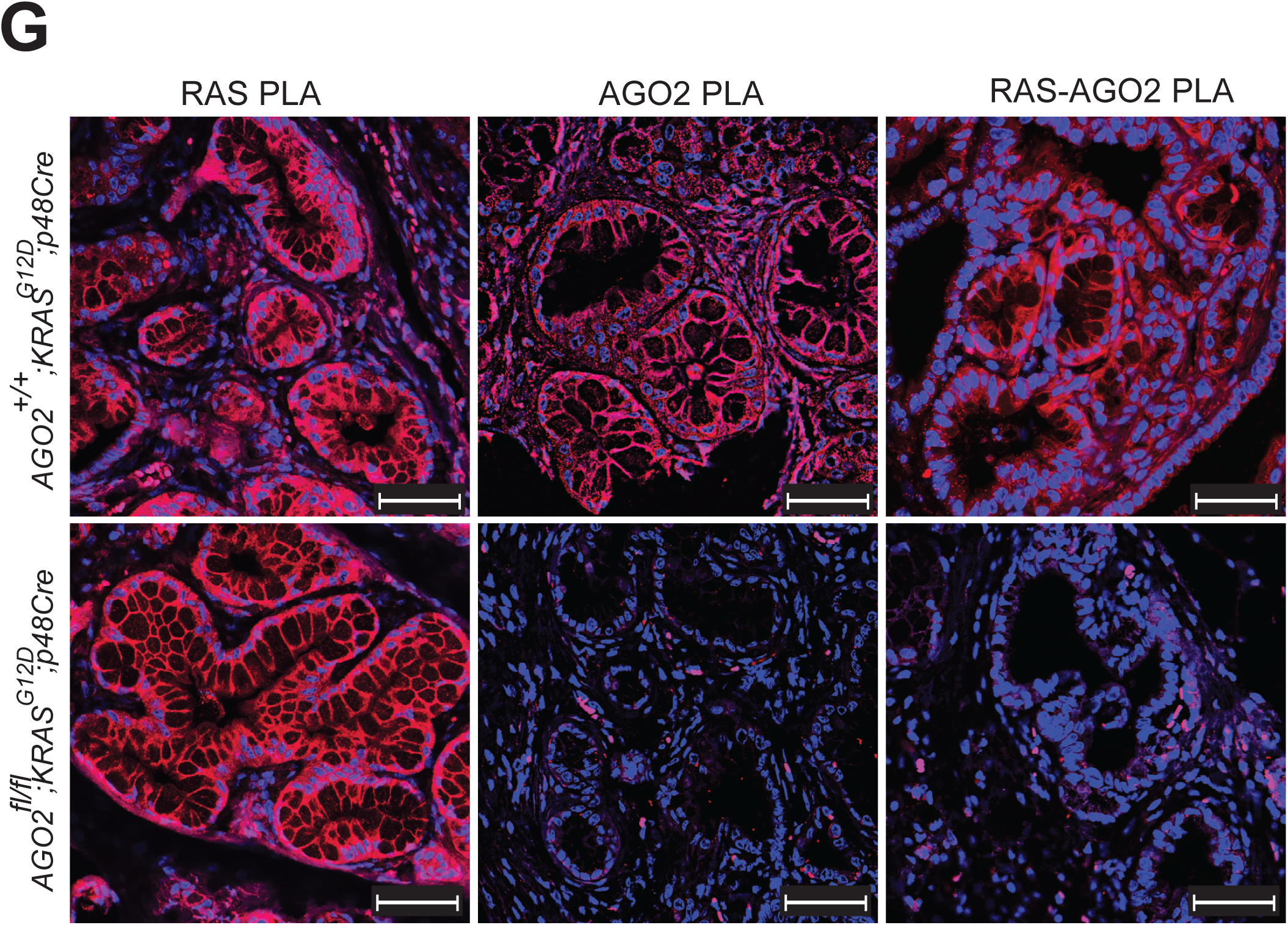
Increased membrane co-localization of RAS and AGO2 during mouse and human PDAC progression. (A) RAS10 (panRAS) antibody specificity for IHC and IF analyses was determined by staining RASless MEFs rescued by either oncogenic *KRAS* or *BRAF*^*V600E*^. Scale bar, 100 µm. (B) Membranous RAS staining in 10-week old PanINs of mouse tissues expressing oncogenic *KRAS* using either IHC (left) or IF (right). Scale bars, 50 µm. (C) Peptide competition assay to demonstrate specificity of the RAS10 antibody in mouse tissues expressing oncogenic *KRAS*. Representative IF images using the RAS10 antibody pre-incubated with RAS peptide spanning the antibody epitope 30-39aa and control overlapping RAS peptide spanning 34-43aa. Scale bar, 50 µm. (D) Representative images of AGO2 IF analysis in pancreatic tissues from *AGO2*^*+/+*^;*KRAS*^*G12D*^;*p48Cre* and *AGO2*^*fl/fl*^;*KRAS*^*G12D*^;*p48Cre* mice. (E) Representative images of IF analysis for RAS and AGO2 through PDAC progression in the *AGO2*^*+/+*^;*KRAS*^*G12D*^;*p48Cre* mice. (F) Representative images of IF analysis of human pancreatic tissue on a TMA showing co-localization of AGO2 and RAS in PanIN and PDAC cells. For (E) and (F), numbers adjacent to merged images indicate the Pearson’s coefficient of co-localization (PCC) of RAS-AGO2 signals at the membranous regions (where 0 is no overlap and 1 is complete overlap). PCC was determined using co-localization signals of at least 50 cells in three distinct areas representative of normal acinar, PanIN, PDAC, or metastases. Scale bar, 50 µm. (G) Representative images of Proximity Ligation Assay (PLA), performed to detect either RAS (RAS PLA) or AGO2 (AGO2 PLA) expression and the RAS-AGO2 interaction (RAS-AGO2 PLA) within PanIN lesions of *AGO2*^*+/+*^;*KRAS*^*G12D*^;*p48Cre* (upper panel) and *AGO2*^*fl/fl*^;*KRAS*^*G12D*^;*p48Cre* (lower panel) mice. PLA signals appear as red dots around DAPI stained nuclei in blue. Scale bar, 50 µm.

### *AGO2* loss restricts progression of PanINs to PDAC through oncogene-induced senescence

Since precancerous lesions have been shown to undergo oncogene-induced senescence (OIS) in the pancreatic cancer mouse model^23^, we postulated that this process may be occurring in the absence of *AGO2*. To test this hypothesis, we performed OIS-associated β-galactosidase staining in pancreatic tissue sections of *AGO2*^*+/+*^;*KRAS*^*G12D*^;*p48Cre* and *AGO2*^*fl/fl*^;*KRAS*^*G12D*^;*p48Cre* mice. As shown in **Fig. 2a-b**, PanINs of *AGO2*^*fl/fl*^;*KRAS*^*G12D*^; *p48Cre* mice showed a significant increase in senescence at the early time point that dramatically increased at 500 days compared to those with *AGO2*-expression. Interestingly, immunoblot analysis of pancreatic tissues obtained from *AGO2*^*fl/fl*^;*KRAS*^*G12D*^;*p48Cre* mice revealed a significant increase in phospho-ERK levels compared to *AGO2*^*+/+*^;*KRAS*^*G12D*^;*p48Cre* mice (which progress to PDAC), indicative of hyperactive MAPK signaling downstream of RAS in the absence of *AGO2* (**Fig. 2c**). This striking observation resembles the effects of oncogenic *BRAF*^*V600E*^ in the pancreas^24^. Indeed, Collisson et. al. showed that *BRAF*^*V600E*^ alone was sufficient to drive PanIN formation that failed to progress to PDAC, despite active MAPK signaling. Consistent with immunoblot analysis, phospho-ERK also showed strong and uniform IHC staining within PanINs in samples with *AGO2* ablation (**Supplementary Fig. 6**). By contrast, phospho-ERK staining was not uniformly detected in the PDACs from *AGO2*^*+/+*^;*KRAS*^*G12D*^;*p48Cre* mice. Thus, oncogenic *KRAS*-driven progression from PanIN to PDAC requires *AGO2* expression to block OIS in mice. We also observed OIS with high levels of phospho-ERK staining in PanINs from human pancreatic tissue (**Fig. 2d**), suggesting that similar mechanisms may block PDAC development in human tissue.

**Figure 6.**
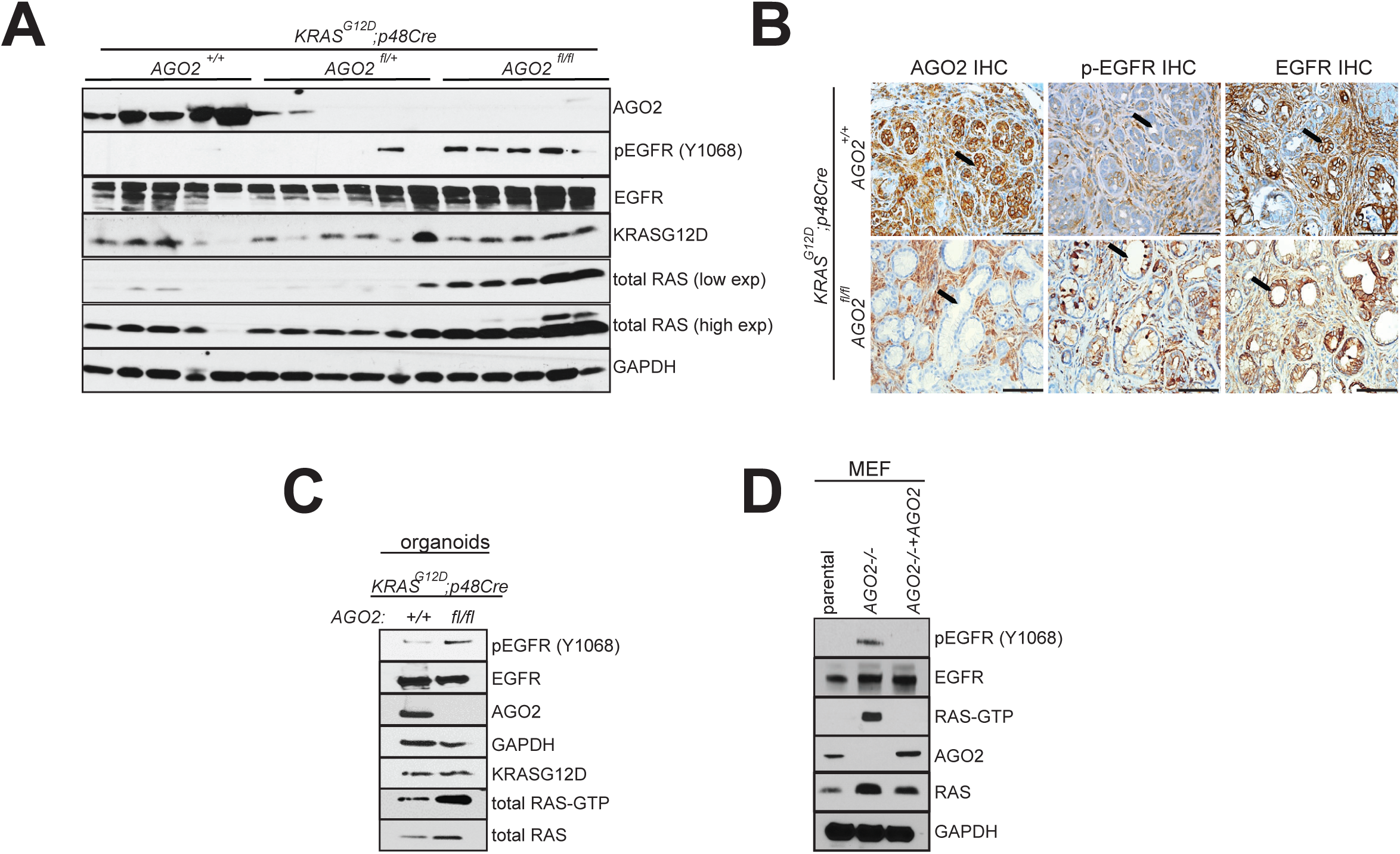
*AGO2* loss activates EGFR-RAS signaling in PanINs. (A) Immunoblot analysis of AGO2 and associated RAS signaling molecules from individual pancreata obtained from 12-week old mice of the indicated genotypes. (B) Representative images of IHC analysis for pEGFR, EGFR, and AGO2 in PanINs of 12-week old mice in the *AGO2*^*+/+*^;*KRAS*^*G12D*^;*p48Cre* and *AGO2*^*fl/fl*^;*KRAS*^*G12D*^;*p48Cre* groups. Arrows indicate PanINs in each panel. Scale bar is 100 µm. (C) Immunoblot analysis of EGFR and wild-type RAS activation levels in pancreatic ductal organoids obtained from 12-week old *AGO2*^*+/+*^;*KRAS*^*G12D*^;*p48Cre* and *AGO2*^*fl/fl*^;*KRAS*^*G12D*^;*p48Cre* mice. Organoids were cultured in the absence of EGF in culture medium for 9 weeks. Total RAS-GTP was determined using the RAF binding assay followed by immunoblotting with pan-RAS antibodies. (D) Immunoblot analysis of EGFR and wild-type RAS activation levels in lysates of parental, *AGO2-/-*, and *AGO2-/-*+ *AGO2* mouse embryonic fibroblasts (MEF). RAS-GTP levels were determined by the RAF binding assay.

### Concomitant oncogenic *KRAS* activation and *p53* loss overcome the requirement for *AGO2* in PDAC progression

Since p53 loss leads to evasion of senescence^25^ and mutational inactivation of the *TP53* gene has been observed in approximately 75% of PDAC patients^26^, we determined the role of AGO2 in the context of p53 loss. For these studies, ablation of *AGO2* expression was carried out in the *Kras*^*G12D*^;*Trp53*^*fl/+*^;*p48Cre* (KPC) mouse model. In these mice, Cre activation simultaneously activates *KRAS* and reduces *Trp53* levels. Unlike the previous mouse model, tumor-free survival of chimeric mice with *AGO2*^*+/+*^;*Kras*^*G12D*^;*Trp53*^*fl/+*^;*p48Cre, AGO2*^*fl/+*^;*Kras*^*G12D*^;*Trp53*^*fl/+*^;*p48Cre*, and *AGO2*^*fl/fl*^;*Kras*^*G12D*^;*Trp53*^*fl/+*^;*p48Cre* genotypes showed no significant differences (**Fig. 3a**). PDAC and metastatic spread were also similar in all of the genotypes analyzed (**Fig. 3b**). IHC confirmed that AGO2 ablation was efficient (**Fig. 3c**). These findings from the two pancreatic mouse models demarcate a role for AGO2 in overcoming OIS to drive PDAC progression. This requirement for *AGO2* can be bypassed through accumulation of *TP53* aberrations^25^, considered a late event in the development of pancreatic cancer^27^.

### Expression and co-localization of RAS and AGO2 in mouse and human pancreatic adenocarcinoma

Having identified an essential role for *AGO2* in PDAC progression in mice, expression levels of AGO2 were next analyzed. Consistent with a role of *AGO2* in *KRAS*-driven oncogenesis in *AGO2*^*+/+*^;*KRAS*^*G12D*^;*p48Cre* mice, IHC analysis showed increased levels of AGO2 in PDAC and metastatic tissues as compared to early PanIN lesions in this model (**Fig. 4a**). To test these observations in human pancreatic cancer, we performed a systematic IHC analysis of a human pancreatic tissue microarray (TMA), comprising 44 duplicate pancreatic tissue cores, including PanIN, PDAC, and metastatic PDAC samples. AGO2 expression was remarkably higher in PDAC and metastatic PDAC cells compared to PanINs (**Fig. 4b**), and this increase was statistically significant (**Fig. 4c**). These data show that AGO2 protein levels are elevated with disease progression and suggest an important role for *AGO2* in pancreatic cancer development in humans.

Considering that active RAS is known to localize to the plasma membrane^2,28^, we tested if RAS and the RAS-AGO2 interaction could be localized at the plasma membrane in the mouse models and human tissues. Since most commercial KRAS-specific antibodies have been shown to be unsuitable for IHC or immunofluorescence (IF)^29^, we tested RAS10, a pan-RAS monoclonal antibody that showed specific staining, as determined by the total loss of signal in RASless MEFs (**Fig. 5a** and **Supplementary Fig. 7a-b**). Surprisingly, relative to the surrounding normal tissue, IHC and IF analysis of mouse pancreatic tissues with this antibody detected high membranous RAS expression within the PanINs (**Fig. 5b**), indicative of activated RAS. Notably, pre-incubation of the antibody with RAS peptides spanning the antibody epitope abrogated RAS staining (**Fig. 5c**), further demonstrating the specificity of the antibody to detect RAS. To corroborate the finding that RAS IHC and IF staining were primarily restricted to oncogenic KRAS-driven PanINs, we performed RNA *in situ* hybridization (RNA-ISH) using *KRAS*-targeted RNA probes (**Supplementary Fig. 7c**). As shown in **Supplementary Fig. 7d**, we observed *KRAS* transcripts restricted to the ducts of pancreatic lesions. The elevated *KRAS* transcript expression in PanINs is consistent with a recent study reporting increased oncogenic *KRAS* transcripts in engineered mouse models^30^. We also validated the AGO2 monoclonal antibody by using pancreatic tissue from *AGO2*^*fl/fl*^;*KRAS*^*G12D*^;*p48Cre* mice (**Fig. 5d** and **Supplementary Fig. 2**).

**Figure 7.**
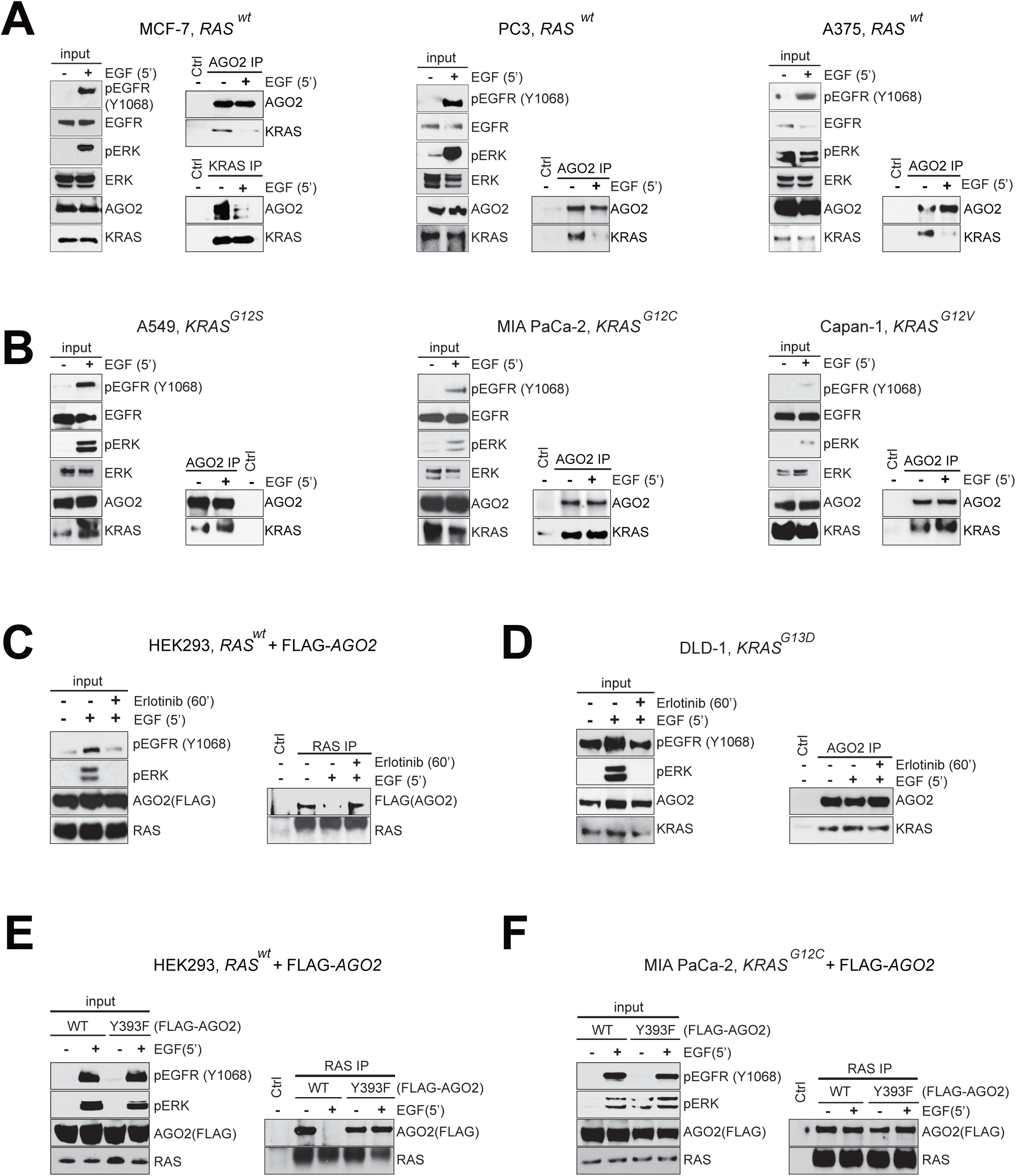
EGFR phosphorylation of AGO2^Y393^ disrupts wild-type KRAS-AGO2 interaction at the membrane but not the mutant KRAS-AGO2 interaction. (A) Immunoprecipitation (IP) of endogenous AGO2 upon EGF stimulation (5’) in the indicated cancer cells expressing wild-type *RAS* followed by immunoblot analysis of KRAS. For MCF7 cells, endogenous co-IP analysis was performed using both AGO2 and KRAS specific antibodies. For each cell line and panel in this figure, MAPK activation and levels of various proteins are shown as input blots. (B) IP of endogenous AGO2 upon EGF stimulation (5’), in the indicated cancer cells harboring different *KRAS* mutations, followed by immunoblot analysis of KRAS. (C) Co-IP and immunoblot analysis of RAS and AGO2 upon EGF stimulation of HEK293 (*KRAS*^*WT*^) cells expressing FLAG-AGO2 or (D) DLD-1 (*KRAS*^*G13D*^) cells in the presence or absence of erlotinib. (E) EGF stimulation and RAS co-IP analysis in HEK293 (*KRAS*^*WT*^) and (F) MIA PaCa-2 (*KRAS*^*G12C*^) cells expressing FLAG-tagged *AGO2* (WT or Y393F). (G) Upper panels, Representative images of single target (RAS or AGO2) and RAS-AGO2 interaction PLA in wild type RAS expressing PC3 cells across the indicated cell culture conditions. Lower panels, Representative images of PLA to detect RAS-AGO2 interaction in wild-type RAS expressing MCF-7 (panel I) and oncogenic KRAS expressing HCT116 (panel II) and H358 (panel III) cells grown in the indicated culture conditions. PLA signals appear as red dots around DAPI stained nuclei in blue. Scale bar, 50 µm.

Having validated the antibodies for detection of RAS and AGO2, we next assessed whether RAS and AGO2 co-localized in pancreatic tissues during PDAC progression. IF staining of normal acinar cells in the mouse pancreas displayed low and diffuse cytoplasmic staining of RAS and minimal expression of AGO2, with a low measure of co-localization (Pearson’s correlation for co-localization, PCC=0.1) (**Fig. 5e**). Interestingly, as shown in **Fig. 5e**, expression of KRAS^G12D^ showed an increased level of activated RAS in PanINs, which further increased in PDAC. In a parallel manner, AGO2 expression progressively increased in PanIN and PDAC tissues with a concomitant increase in plasma membrane localization and co-localization with RAS (PanIN PCC=0.5; PDAC PCC=0.7). Localization of AGO2 to the plasma membrane was independently confirmed by analyzing co-localization with the membrane marker, E-cadherin (PCC=0.43; **Supplementary Fig. 8a**). Furthermore, in pancreatic tissue obtained from normal mice treated with caerulein to induce pancreatitis^31^, we observed wild-type RAS localization at the membrane without AGO2 co-localization (**Supplementary Fig. 8b**). This suggests specificity of the RAS-AGO2 co-localization observed during oncogenic KRAS-driven PDAC development. IF analyses continued to exhibit significant membranous co-localization signals for RAS and AGO2 (PCC=0.7) in PDAC lesions from *AGO2*^*+/+*^;*Kras*^*G12D*^;*Trp53*^*fl/+*^;*p48Cre* mice (**Supplementary Fig. 8c**). Importantly, extending the IF analysis to human pancreatic tissues, we observed a similar pattern of localization of RAS and AGO2 with increased RAS-AGO2 co-localization signals at the plasma membrane associated with pancreatic cancer progression (PCC, normal to PDAC increased from 0.1 to 0.5, respectively) (**Fig. 5f**).

**Figure 8.**
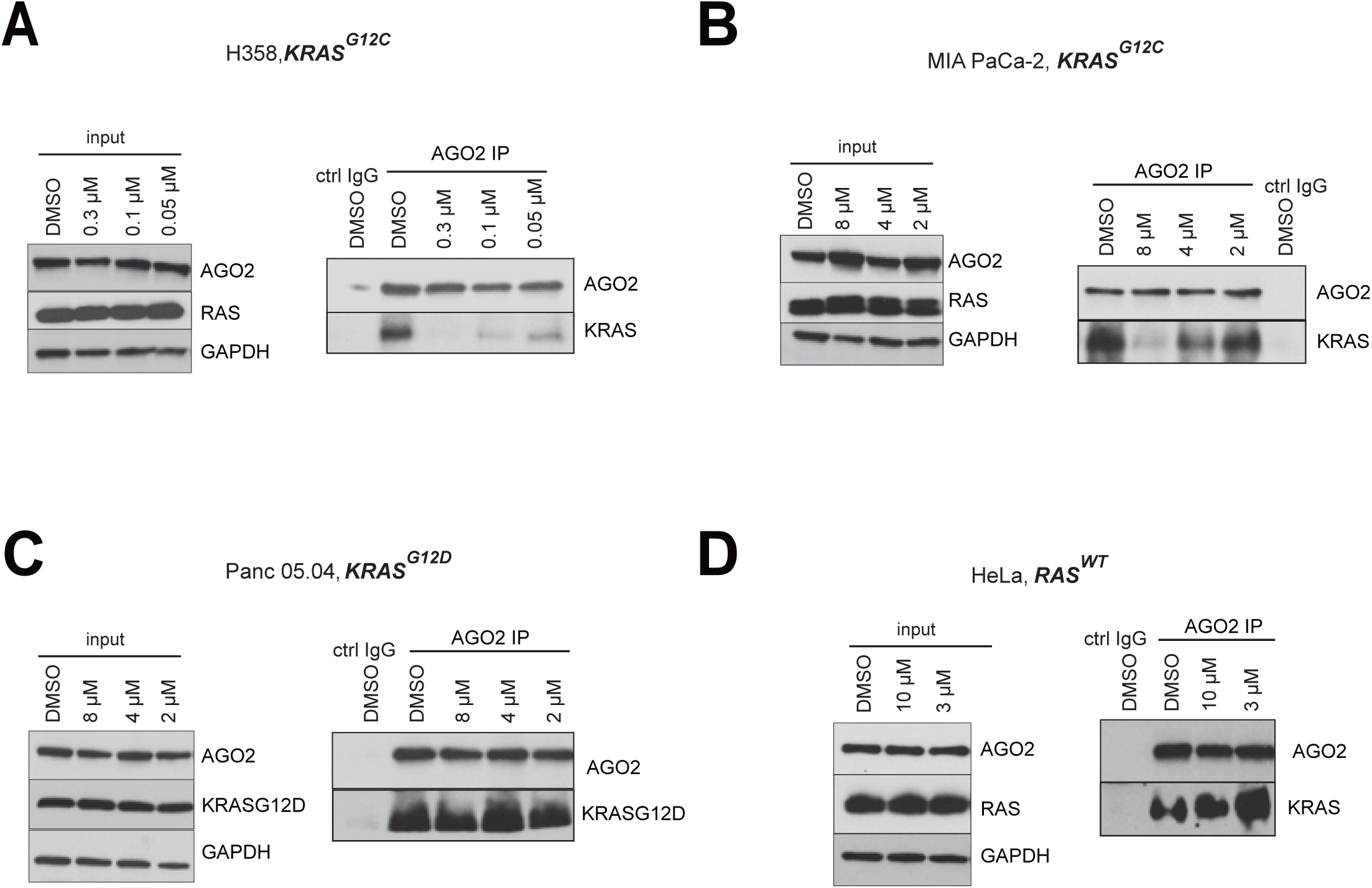
ARS-1620, a G12C-specific inhibitor, disrupts the KRAS^G12C^-AGO2 interaction. IP of endogenous AGO2 followed by IB to detect KRAS, in *KRAS*^*G12C*^ harboring (A) H358 and (B) MIA PaCa-2 cells treated with varying concentrations of ARS-1620 for 3 and 9 hours, respectively. *KRAS*^*G12D*^ and *KRAS*^*wt*^ harboring (C) Panc 05.04 and (D) HeLa cells, respectively, treated with ARS-1620 for 24h followed by AGO2 IP and immunoblot analysis of KRAS^G12D^ or KRAS. For each cell line, input blots for AGO2 and RAS are shown.

For a direct assessment of the RAS-AGO2 interaction at single molecule resolution, we performed proximity ligation assays (PLA)^32,33^ using the RAS and AGO2 antibodies validated earlier (**Fig. 5a** and **Supplementary Fig. 2, 8d**). As shown in **Fig. 5g**, PLA signals indicative of RAS-AGO2 interaction were observed at the plasma membrane within PanINs arising in *AGO2*^*+/+*^;*KRAS*^*G12D*^;*p48Cre* but not in *AGO2*^*fl/fl*^;*KRAS*^*G12D*^;*p48Cre* mice. This further corroborates the IF analyses and provides the first evidence of membranous RAS-AGO2 interaction. Together these data indicate that during pancreatic cancer development, AGO2 localizes at the plasma membrane, the site of RAS activity^2,28^, and substantiates a role for *AGO2* in the progression of PanINs to PDAC.

### Loss of *AGO2* and its impact on the EGFR-RAS signaling axis

The data above place AGO2 and RAS together at the plasma membrane during PDAC progression. Therefore, we next sought to explore how AGO2 may mediate alteration of RAS signaling pathways. We specifically focused on the EGFR-RAS signaling axis for two reasons: 1) EGFR has been shown to be essential for PanIN formation in the *KRAS*^*G12D*^-driven pancreatic mouse model that we have used^10,11,34^, and 2) EGFR activation has been shown to directly inhibit AGO2 function through phosphorylation of its tyrosine 393 residue^16^. Immunoblot analysis of pancreatic tissues from 12-week old mice with *AGO2*^*+/+*^;*KRAS*^*G12D*^;*p48Cre, AGO2*^*fl/+*^;*KRAS*^*G12D*^;*p48Cre*, and *AGO2*^*fl/fl*^;*KRAS*^*G12D*^;*p48Cre* genotypes showed a marked increase in AGO2 levels relative to normal pancreata (**Supplementary Fig. 9**) concordant with IF analysis (**Fig. 5e**). Also, consistent with published studies^10,11^, total EGFR levels were elevated in *KRAS*^*G12D*^ mice irrespective of the *AGO2* genotype (**Supplementary Fig. 9**). However, in early PanINs initiated by oncogenic KRAS, significantly higher levels of phospho-EGFR (Y1068) were observed in pancreatic tissues of *AGO2*^*fl/fl*^;*KRAS*^*G12D*^;*p48Cre* mice (**Fig. 6a** and **Supplementary Fig. 9**), indicating activated EGFR signaling in the absence of *AGO2* expression. IHC analysis confirmed that the elevated phospho-EGFR levels observed in tissue lysates were restricted to the PanIN lesions of *AGO2*^*fl/fl*^;*KRAS*^*G12D*^;*p48Cre* mice (**Fig. 6b**). IHC of total EGFR showed no significant difference in expression in pancreatic tissues between the two genotypes (**Fig. 6b** and **Supplementary Fig. 10a**). As previously noted, irrespective of *AGO2* genotype, lesions from later time points showed a marked reduction in total EGFR levels^10,11^ in mouse (and with disease progression in human tissue), further supporting the significance of EGFR signaling in the early stages of disease (**Supplementary Fig. 10a-b**). Importantly, immunoblot analysis showed that EGFR activation was accompanied with a remarkable increase in total RAS levels but not oncogenic KRAS^G12D^ levels (**Fig. 6a** and **Supplementary Fig. 9**), raising an intriguing possibility that growth factor activation involves signaling along the EGFR-wild-type RAS axis in early stage PanINs.

**Figure 9.**
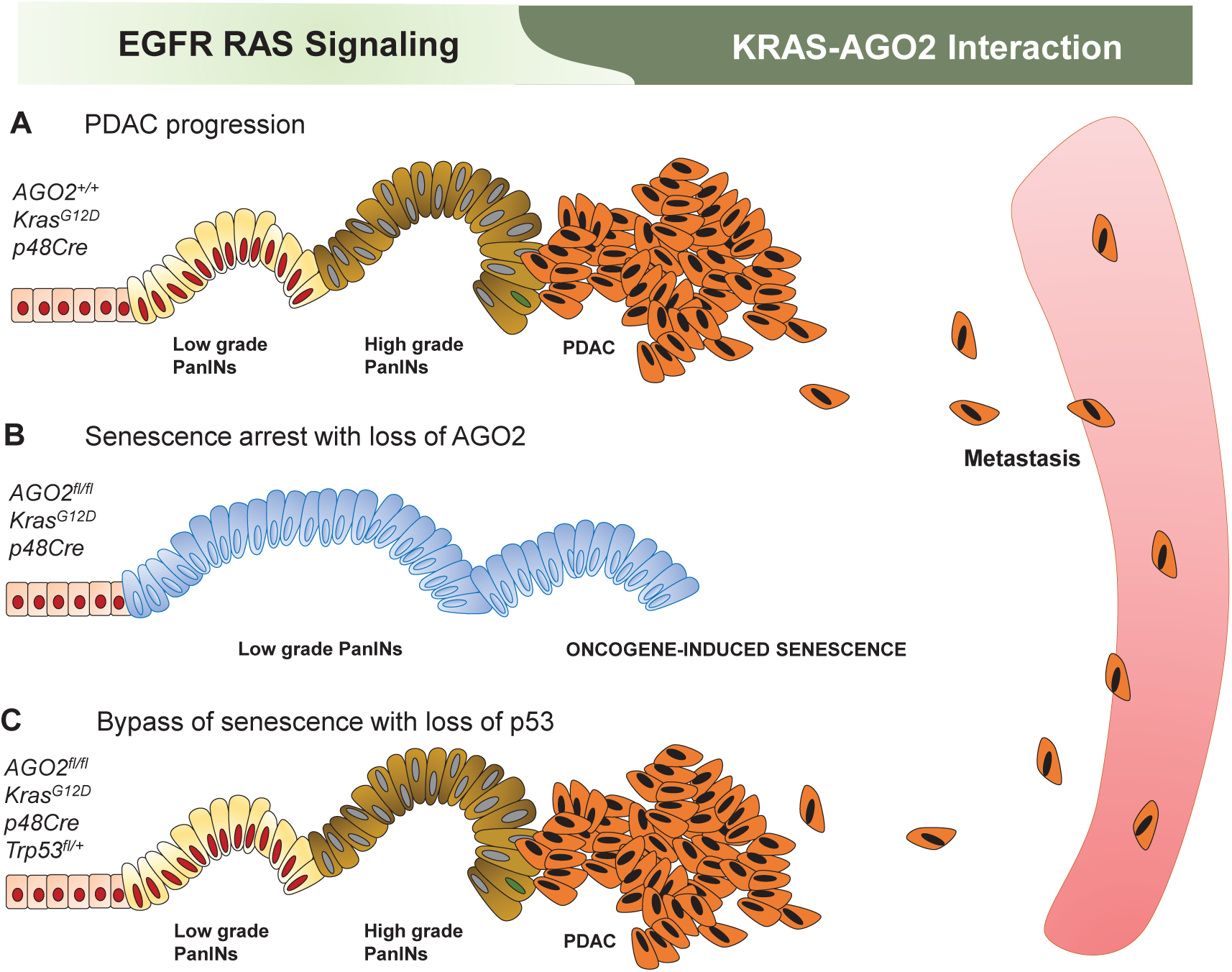
Schematic model showing the essential role of *AGO2* in pancreatic cancer progression. (A) Expression of *KRAS*^*G12D*^ in normal pancreatic cells initiates low grade PanINs which progress to higher grade PanINs, PDAC, and metastases. (B) *AGO2* loss leads to oncogene-induced senescence and prevents progression of low grade PanINs to PDAC. (C) Loss of *p53* in this model leads to bypass of senescence mechanisms, and progression from PanIN to PDAC is independent of AGO2. The study supports an early phase of PanIN formation which is EGFR/RAS signaling dependent, followed by a dependency on the RAS-AGO2 interaction for PDAC progression.

To investigate this further, we isolated pancreatic ducts from 12-week old *AGO2*^*+/+*^;*KRAS*^*G12D*^;*p48Cre* and *AGO2*^*fl/fl*^;*KRAS*^*G12D*^;*p48Cre* mice and cultured them as organoids^35^ in the absence of EGF (**Supplementary Fig. 11**). Immunoblot analysis showed increased levels of phospho-EGFR and total RAS in the organoids with *AGO2* loss, while KRAS^G12D^ expression showed no change (**Fig. 6c**), mirroring the observations from pancreatic tissue lysates. Remarkably, the total RAS levels (which includes both wild-type and mutant RAS), as well as total RAS-GTP, were higher in organoids deficient in *AGO2*.

To probe if *AGO2* loss activates wild-type RAS even in the absence of mutant KRAS, we performed immunoblot analysis and RAS activation assays using *AGO2*^-/-^ MEFs that do not harbor any form of oncogenic RAS^36^. As shown in **Fig. 6d**, *AGO2*^*-/-*^ MEFs also exhibit increased phospho-EGFR and wild-type RAS levels along with elevated wild-type RAS-GTP levels, which were significantly reduced when rescued with *AGO2*. Given that AGO2 is a direct phosphorylation substrate of the EGFR kinase^16^, our experiments define a previously unknown reverse feedback up-regulation of phospho-EGFR via AGO2 that controls RAS activation^37-40^.

### AGO2^Y393^ phosphorylation by EGFR disrupts wild-type, but not mutant, KRAS-AGO2 interaction at the membrane

Considering that the loss of *AGO2* expression leads to activation of both EGFR and wild-type RAS (**Fig. 6**), we posited that AGO2 binding to KRAS may represent a rate limiting step in the activation of wild-type KRAS during growth factor stimulation. To explore this premise, we assayed for KRAS-AGO2 interaction across a panel of cell lines expressing wild-type or mutant *RAS* stimulated with EGF. Interestingly, EGF stimulation resulted in a dramatic decline in KRAS-AGO2 interaction in cells with wild-type *KRAS*, as seen in MCF-7, PC3, A375, and HeLa cells (**Fig. 7a** and **Supplementary Fig. 12a-b**). In stark contrast, EGF stimulation of cells harboring oncogenic *KRAS*, including A549 (*KRAS*^*G12S*^), MIA PaCa-2 (*KRAS*^*G12C*^), and Capan-1 (*KRAS*^*G12V*^), retained binding of endogenous KRAS and AGO2 despite activation of the EGFR/MAPK/AKT pathway (**Fig. 7b** and **Supplementary Fig. 12c**). Disruption of the wild-type RAS-AGO2 interaction was also observed when HEK293 (*KRAS*^*WT*^) cells expressing FLAG-tagged AGO2 were stimulated with EGF; the interaction was rescued by treatment of cells with the EGFR kinase inhibitor, erlotinib (**Fig. 7c**). This strongly suggests that EGFR kinase activity was critical for the disruption of the wild-type KRAS-AGO2 interaction. In contrast, DLD-1 cells harboring mutant *KRAS*^*G13D*^ showed no loss of KRAS and AGO2 association either by EGF treatment, or by EGFR kinase inhibition with erlotinib (**Fig. 7d**).

To test if the previously identified site of EGFR-mediated phosphorylation^16^ on AGO2 at tyrosine 393 has a role in binding to KRAS, we analyzed the ability of a phosphorylation-deficient AGO2^Y393F^ mutant to bind RAS under different conditions. In HEK293 (*KRAS*^*WT*^) cells, EGF stimulation led to dissociation of wild-type AGO2 from RAS, but the AGO2^Y393F^ mutant continued to bind RAS with or without EGFR activation (**Fig. 7e**), indicating that phosphorylation of this residue is critical for dissociation. Expression of these *AGO2* constructs in MIA PaCa-2 (*KRAS*^*G12S*^) cells showed no discernible change in RAS binding upon EGFR activation (**Fig. 7f**).

To track the localization of the RAS-AGO2 interaction upon growth factor activation, we performed PLA on cells expressing wild-type or mutant *KRAS*. The use of either RAS or AGO2 antibodies alone did not show signals for the RAS-AGO2 PLA (**Supplementary Fig. 13a**). Interestingly, serum starved PC3 cells showed increased membrane localization of both RAS and AGO2 proteins contributing to the dramatic increase in membrane localized RAS-AGO2 PLA signals under these conditions (**Fig. 7g**, upper panels). RAS-AGO2 interaction PLA signals were significantly reduced upon EGF stimulation and restored to levels observed under serum sufficient conditions. IF analyses also showed a similar pattern of RAS-AGO2 co-localization under these different culture conditions (**Supplementary Fig. 13b**). A similar pattern of RAS-AGO2 interaction PLA signals was observed in wild-type *RAS* expressing MCF-7 cells (**Fig. 7g**, panel I). In contrast, both HCT116 and H358 cells (**Fig. 7g**, panels II and III), expressing oncogenic forms of *KRAS*, showed higher basal levels of RAS-AGO2 PLA signals compared to wild-type *RAS* expressing cells that remained consistent under different cell culture conditions. This further corroborates results obtained above through co-IP analysis. Combined, these data suggest that the wild-type KRAS-AGO2 interaction at the membrane is sensitive to EGFR-mediated phosphorylation of AGO2^Y393^, while the oncogenic KRAS-AGO2 interaction is unaffected by both EGFR activation status as well as AGO2^Y393^ phosphorylation.

### ARS-1620, a G12C inhibitor, disrupts the oncogenic KRAS-AGO2 interaction

Finally, we tested if direct targeting of oncogenic KRAS could affect the endogenous mutant KRAS-AGO2 interaction. Interestingly, the mutant KRAS-AGO2 interaction was disrupted when H358 (**Fig. 8a**) and MIA-PaCa-2 cells (**Fig. 8b**), harboring *KRAS*^*G12C*^ mutant alleles, were treated with ARS-1620^41^, a covalent G12C inhibitor. The disruption of endogenous KRAS^G12C^-AGO2 interaction in these cells was concentration dependent and reflects the differential sensitivities of the two cell lines to ARS-1620^42^. In a similar assay, ARS-1620 treatment had no effect on the KRAS^G12D^-AGO2 or KRAS^wt^-AGO2 interaction in Panc 05.04 cells (**Fig. 8c**) or HeLa cells (**Fig. 8d**), respectively. Given that ARS-1620 binds an allosteric Switch II pocket (SW-IIP)^42^ on GDP-loaded KRAS^G12C^, the disruption of the KRAS^G12C^-AGO2 binding provides orthogonal evidence that AGO2 makes contact with the Switch II region in KRAS. This data also proves that besides SOS, the easily detectable, endogenous membrane bound KRAS^G12C^-AGO2 interaction (**Fig.7g**) is an additional target of G12C inhibitors.

## DISCUSSION

Genetically engineered mouse models have been extensively used to mirror the stepwise progression of human pancreatic cancer, starting with benign precursor lesions (PanINs) driven by mutant *KRAS*^9,43,44^. Here, using GEMM models of *AGO2* loss, we propose that pancreatic cancer development involves two phases (**Fig. 9a**). In the first phase, PanIN development is triggered by oncogenic *KRAS* and depends on EGFR-RAS mediated proliferation, where persistent signaling along this axis can initiate OIS. While low-grade PanINs have been known to undergo senescence in pancreatic cancer mouse models^23^, our model with *AGO2* loss represents the first instance where OIS completely abrogates the progression of PanIN to PDAC (**Fig. 9b**). Failure to detect bypass routes to oncogenesis, unlike the GEMM model with *NOTCH2* loss^45^, highlights the robustness of OIS observed in the absence of *AGO2*. The second phase of PDAC progression is AGO2 dependent, and the AGO2-KRAS interaction likely plays a predominant role (**Fig. 9a**). AGO2 is found localized at the plasma membrane, a site not previously associated with RNA silencing activity^46-49^. Importantly, we show for the first time that AGO2 levels are increased in PDAC and metastases compared to PanIN lesions from patients, highlighting the relevance of AGO2 in clinical disease progression. Increased RAS-AGO2 co-localization at the plasma membrane is also seen during PDAC progression in mice and humans, providing evidence for the role of AGO2 at the site of oncogenic KRAS activity. We hypothesize that this robust interaction between mutant KRAS and AGO2 overcomes the OIS block, facilitating the second phase of progression, from PanIN to PDAC. Loss of *p53* precludes this requirement for *AGO2* (**Fig. 9c**), but RAS-AGO2 co-localization is retained in this model.

In addition to delineating a biphasic model of pancreatic cancer development, this study defines novel aspects of the EGFR-RAS-AGO2 signaling network, particularly regarding modulation of the RAS-AGO2 interaction. PLA provides the first evidence of direct RAS-AGO2 interaction at the plasma membrane within pancreatic lesions driven by oncogenic *KRAS*. Interestingly, wild-type *RAS* expressing cells undergoing serum starvation also show increased RAS-AGO2 interaction at the membrane. This suggests that the membrane RAS-AGO2 association under conditions of stress (starvation or presence of oncogenic KRAS) allows fine-tuning of RAS signaling through growth factor receptor activation. Phosphorylation of AGO2 by EGFR simultaneously inhibits the last step of microRNA biogenesis^16^ and activates RAS at the plasma membrane^50^, a previously unrecognized aspect of EGFR-RAS-MAPK signaling. However, this observation is consistent with previously reported disruptions of other protein-protein interactions through EGFR phosphorylation as a strategy to modulate RAS activation signals at the membrane^51-53^.

Our observation that EGF stimulation disrupts the wild-type KRAS-AGO2 interaction, but not the oncogenic KRAS-AGO2 interaction, is likely central to the critical requirement of EGFR signaling in PanIN formation^10,11^. It is also intriguing that EGFR-mediated phosphorylation of AGO2^Y393^ disrupts wild-type RAS binding in a manner reminiscent of AGO2-Dicer binding^16^. However, AGO2-KRAS mutant binding remains unaffected by AGO2 phosphorylation status, underscoring the critical reliance of PDAC progression on this interaction. Notably, we find that a G12C covalent inhibitor can disrupt the mutant KRAS-AGO2 interaction. Altogether, our data suggest that abrogation of the oncogenic KRAS-AGO2 association at the plasma membrane may represent a novel therapeutic opportunity for pancreatic cancer treatment that warrants further investigation.

## Supporting information

All Supplemental Figs

## ACKNOWLEDGMENTS

We thank Mandy Davis and Marta Hernadi-Muller for their help with processing paraffin-embedded slides. We acknowledge Sisi Gao and Stephanie Ellison for their help in manuscript preparation. We also thank Saravana Dhanasekaran and Markus Eberl for technical assistance and Eric Fearon for discussions. S.S. is a Genentech Fellow. A.M.C is an American Cancer Society Research Professor and Taubman Scholar.

## AUTHOR CONTRIBUTIONS

Mouse experimental data were generated by J.C.T., S.C., K.M.J., A.G., A.X., G.T, V.L.D, and S.S. Contributions to other experimental data were made by S.S., R.F.S, V.L.D., S.Z.-W., S.E., Y.Z., M.M., Javed S., I.J.A., and C.K.-S. L.W. and Jiaqi. S. coordinated the pathology assessment. Jiaqi. S. provided the human TMA and performed IHC scoring. X.W. performed RNA ISH. J.W., R.S., H.C, and J.T. performed molecular dynamic simulation. X.C. helped with project management. H.C.C. supervised the pancreatitis experiments (Grants, NIH U01 CA224145 and DOD CA170568). S.S. and A.M.C. jointly conceived the study. S.S., C.K-S., and A.M.C. wrote the manuscript. Funding and overall supervision of the study was provided by A.M.C.

## METHODS

### Mouse strains

LSL-*KRAS*^*G12D*^ (ref. ^6^) (Kras ^LSL-*G12D*^) and *p48Cre mice*^54^ were obtained from Marina Pasca di Magliano, University of Michigan. Conditionally floxed *AGO2* (ref.^17^) (*AGO2*^fl/fl^) mice and *p53*^*fl/+*^ mice were purchased from Jackson labs (Bar Harbor, Maine). PCR genotyping for *KRAS*^*G12D*^;*p48Cre, p53*^*fl/+*^, and *AGO2* alleles, from DNA isolated from mouse tails, was performed using standard methodology. To generate experimental and control mice, *AGO2*^*fl/fl*^ *p48Cre*, and *KRAS*^*G12D*^ lines were intercrossed to generate *AGO2*^*fll+*^;*p48Cre* and *KRAS*^*G12D*^;*p48Cre* mice. These two lines were then intercrossed to generate the *AGO2*^*fllfl*^;*KRAS*^*G12D*^;*p48Cre* experimental mice. Given that mice were maintained on a mixed background, littermate controls were systematically used in all experiments (sex ratio per cohort was balanced). All animals were housed in a pathogen-free environment, and all procedures were performed in accordance with requirements of the University of Michigan IACUC.

Cre activation in acinar cells of pancreata of mice with mutant KRAS alleles was validated by genotyping using the *KRAS*^*G12D*^ conditional PCR primers 5’ *gtc ttt ccc cag cac agt gc* 3’, 5’ *ctc ttg cct acg cca cca gct c* 3’, and 5’ *agc tag cca cca tgg ctt gag taa gtc tgc a* 3’ according to Tyler Jacks lab protocol (https://jacks-lab.mit.edu/protocols/genotyping/kras_cond).

### Histology, immunohistochemistry, and immunofluorescence

Paraffin-embedded tissues from mice were processed using standard methodology. Details of the primary antibodies used for IHC are provided in **Supplementary Table 1**. Immunohistochemistry and immunofluorescence staining were performed using standard techniques. For immunofluorescence, slides were viewed using a Nikon 1A-B confocal microscope. To estimate co-localization of proteins, the Coloc2 program (ImageJ) was used to determine Pearsons coefficient. Cells within the PanIN/PDAC or metastatic regions (from mouse and human tissues), excluding the stromal compartment, were used to determine the extent of overlap. In panels with normal tissue shown in **Fig. 5**, acinar cells were used for co-localization analyses. Average values over three different areas are shown.

### Proximity Ligation Assay (PLA)

Cell lines were cultured in 8-well chamber slides. After the indicated treatment/stimulation, cells were fixed with 4% paraformaldehyde and then permeabilized using 0.1%Tween. Subsequent PLA staining was performed as per the protocol provided by the manufacturer (DUOlink kit, Millipore/Sigma). Mouse RAS10 and rabbit AGO2 antibodies, validated in this study, were used at 1:250 dilution to detect signals either alone or in combination. Negative controls were performed using either single antibody (**Supplementary Fig. 13a**), Rasless MEFs (**Supplementary Fig. 8d**), or tissue lacking AGO2 (**Fig. 5d**). Images were obtained using the Nikon A1B inverted confocal microscope. For mouse tissue PLA, the paraffin-embedded sections were processed as for IF analysis. PLA was then performed using RAS10 or AGO2 antibodies, either alone or in combination, and imaged using the Nikon A1B confocal microscope.

### Human TMA analysis

Pancreatic TMAs and frozen human tissue repositories were established by a pathologist (J.S.) and developed at the Tissue and Molecular Pathology Core in the Department of Pathology, University of Michigan, after IRB approval. IHC scoring was performed by a pathologist (J.S.).

### RNA *in situ* Hybridization (RNA-ISH)

RNA-ISH was performed to detect *Kras* mRNA on formalin-fixed paraffin-embedded (FFPE) tissue sections using the RNAscope 2.5 HD Brown kit (Advanced Cell Diagnostics, Newark, CA) and target probes against mouse *Kras* (412491). *Mm-Ubc* (mouse ubiquitin C) and *DapB* (Bacillus bacterial dihydrodipicolinate reductase) were used as positive and negative controls, respectively. FFPE tissue sections were baked for 1 hour at 60°C, deparaffinized in xylene twice for 5 minutes each, and dehydrated in 100% ethanol twice for 1 minute each, followed by air drying for 5 minutes. After hydrogen peroxide pre-treatment and target retrieval, tissue samples were permeabilized using Protease Plus and hybridized with the target probe in the HybEZ oven for 2 hours at 40°C. After two washes, the samples were processed for a series of signal amplification steps. Chromogenic detection was performed using DAB, counterstained with 50% Gill’s Hematoxylin I (Fisher Scientific, Rochester, NY).

### Quantitative RT-PCR

Pancreatic total RNA was isolated using the AllPrep DNA/RNA/miRNA Universal Kit (Qiagen). For quantitation of mRNA transcripts, RNA was extracted from the indicated samples, and cDNA was synthesized using the SuperScript III System according to the manufacturer’s instructions (Invitrogen). Quantitative RT-PCR was conducted using primers detailed in **Supplementary Table 3** with SYBR Green Master Mix (Applied Biosystems) on the StepOne Real-Time PCR System (Applied Biosystems). Relative mRNA levels of the transcripts were normalized to the expression of the housekeeping gene *GAPDH*.

### Pancreatic tissue lysates and immunoblot analysis

Pancreata obtained from mice were homogenized in Mg^2+^-containing lysis buffer. Clear lysates were separated using SDS-PAGE and processed for immunoblot analysis using standard methods. Primary antibodies used in the study are indicated in **Supplementary Table 1**. Particularly, Ras antibodies validated in a recent study^29^ are also indicated.

### Isolation of pancreatic ductal organoids

Pancreatic ductal organoids obtained from 12-week old *KRAS*^*G12D*^;*p48Cre* and *AGO2*^*fl/fl*^;*KRAS*^*G12D*^;*p48Cre* mice were cultured in organoid media as described previously^35^. Organoids were cultured without EGF for 9 passages to exclude normal duct contamination, which are dependent on EGF.

### RAS-GTP analysis

Protein lysates were prepared from pancreatic ductal organoids or cell lines using Mg^2+^-containing lysis buffer. RAF1 binding was used as a measure of RAS-GTP levels in the lysates, as previously described^15^.

### Plasmids

Full length FH-*AGO2* constructs were obtained from Addgene (pIRESneo-FLAG/HA-AGO2 10822, PI:Thomas Tuschl). *AGO2*^*Y393F*^ mutant construct was generated using the QuikChange II XL Site-Directed Mutagenesis Kit (Agilent) from the FH-*AGO2* plasmid described above using the primers hAGO2_Y393F_Fwd 5’*AAATTCACGGACGAATGGATCTGTGTTGAAACTTGCAC*3’ and hAGO2_Y393F_Rev 5’*GTGCAAGTTTCAACACAGATCCATTCGTCCGTGAATTT*3’. DNA sequences were confirmed using Sanger sequencing at the University of Michigan Sequencing Core.

### Cell culture, transfection, and EGF stimulation

All cell lines (detailed in **Supplementary Table 4**) were obtained from the American Type Culture Collection (ATCC). Cells were cultured following ATCC culture methods in media supplemented with the corresponding serum and antibiotics. Additionally, cells were routinely genotyped and tested bi-weekly for mycoplasma contamination. For EGF stimulation, cells were grown to approximately 80% confluence and washed with PBS three times. Cells were incubated overnight (16 hr) in serum free media. EGF stimulation was performed for 5 minutes with 100 ng/µl of epidermal growth factor (Gibco) at 37°C. After stimulation, cells were washed and protein lysates were prepared in K Buffer lysis buffer. For tyrosine kinase inhibition, cells were pre-treated with 15 µM of Erlotinib for 1 hour prior to EGF stimulation, as described above. HEK293 or MIA PaCa-2 cells were transfected with different *AGO2* constructs using Fugene HD (Promega) or Lipofectamine 3000 (Invitrogen) according to the manufacturer’s protocols. For EGFR stimulation with transient *AGO2* construct overexpression, cells were transfected approximately 16 hours prior to overnight serum starvation and EGF stimulation.

RASless MEFs were a kind gift from the RAS Initiative. Details of how these cells were developed and their growth characteristics can be found at https://www.cancer.gov/research/key-initiatives/ras/ras-central/blog/2017/rasless-mefs-drug-screens.

### Immunoprecipitation (IP) Analysis

For immunoprecipitation analysis, protein lysates were prepared in K Buffer (20 mM Tris pH 7.0, 5 mM EDTA, 150 mM NaCl, 1% Triton X100, 1 mM DTT, phosphatase inhibitors, and protease inhibitors). Typically,150-200 µg of protein lysates (RAS10 IP: 150 µg; AGO2 IP: 200 µg; KRAS IP: 150 µg) were pre-cleared with 10 µl of Protein A/G agarose beads (Santa Cruz) for 1 hour. Pre-cleared lysates were incubated with 5-10 µg of the indicated primary antibodies targeting the protein of interest or with corresponding isotype controls overnight at 4°C. 30 µl of Protein A/G beads were then added to immune complexes and incubated for 1-3 hours at 4°C, spun, and washed in 150-300 mM NaCl containing K-buffer prior to separation of immunoprecipitates by SDS-PAGE. To determine the varying levels of KRAS expressed in different cells lines (with or without EGF stimulation), shown in **Fig. 7**., pan RAS10 antibody was used for immunoprecipitation followed by immunoblot analysis using KRAS specific SC-30 antibody.

### β-galactosidase assay

β-galactosidase staining was performed using the Senescence β-Galactosidase Staining Kit #9860 (Cell Signaling) on 10 µM-thick frozen sections of mouse pancreas, as per the manufacturer’s protocol.

