## Supplementary material for "An Essential Role for *Argonaute 2* in EGFR-KRAS Signaling in Pancreatic Cancer Development": All Supplemental Figs

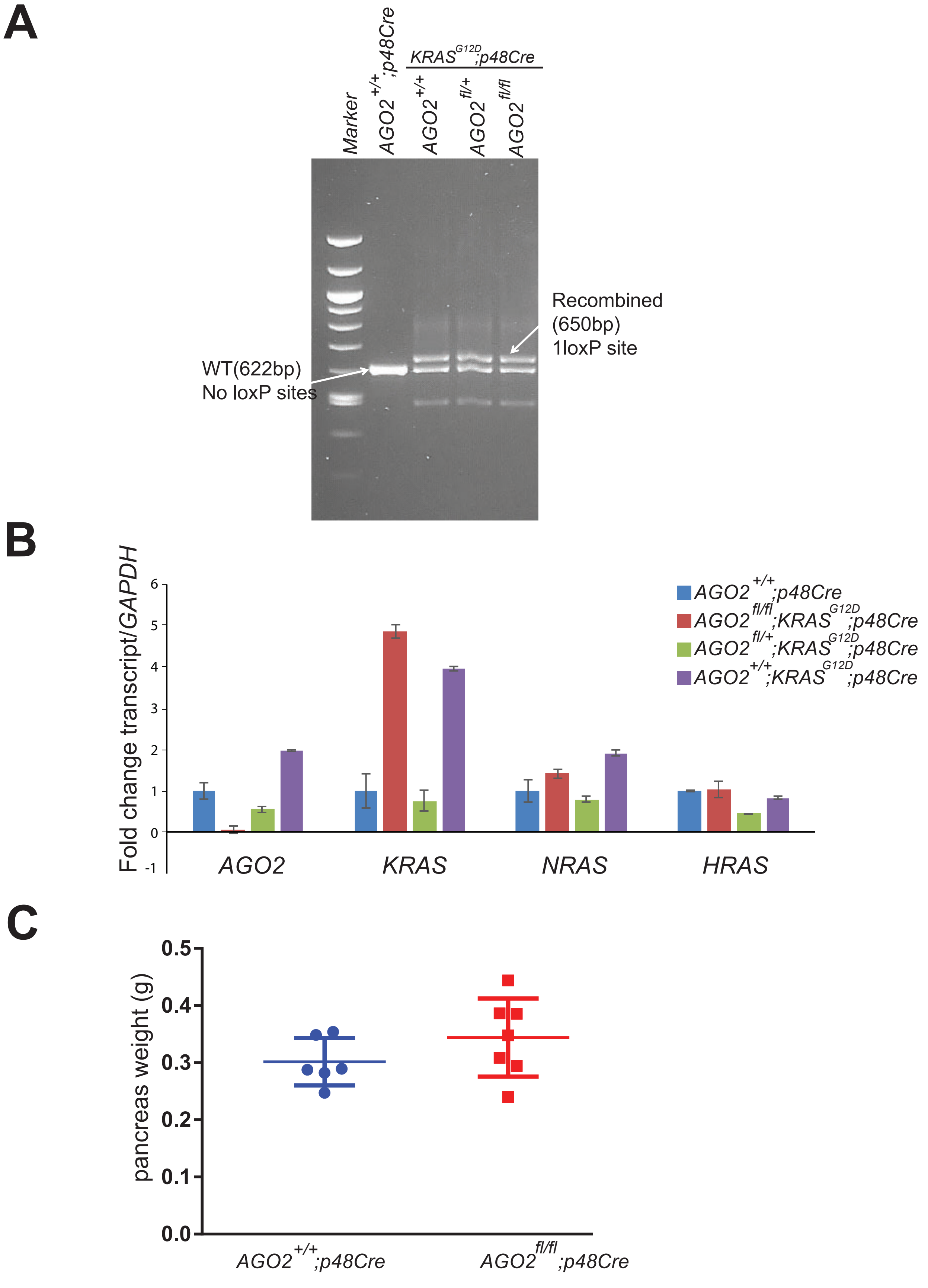


**Supplementary Figure 1: Cre activation in the pancreatic cancer mouse model.** (A) Genomic DNA analysis from the pancreata of the indicated genotypes showing recombined alleles in the LSL-*KRAS* model, as previously described[1]. (B) RT-qPCR analysis for different transcripts from 10-week old mouse pancreata of the indicated genotypes and (C) pancreas weights.

1. Tuveson, D.A., *et al.* Endogenous oncogenic K-ras(G12D) stimulates proliferation and widespread neoplastic and developmental defects. *Cancer Cell* **5**, 375-387 (2004).

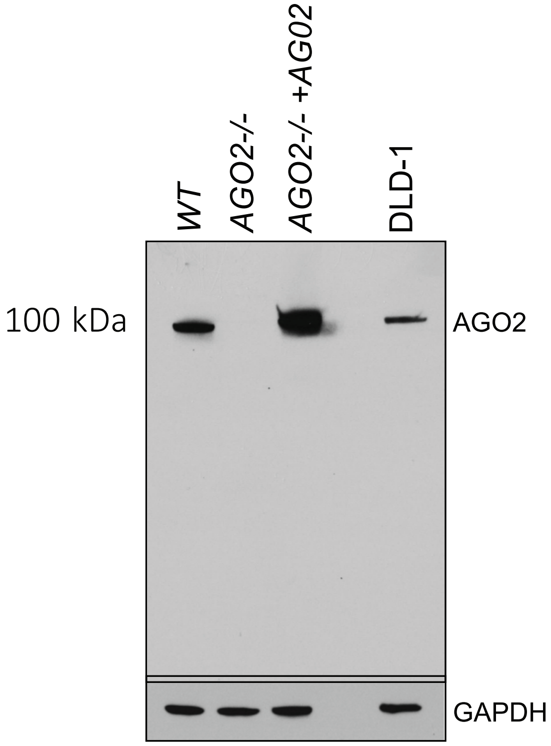


**Supplementary Figure 2: Specificity of AGO2 monoclonal antibody.** Western blot analysis of AGO2-/- mouse embryonic fibroblasts and human colon cancer cells using a monoclonal antibody to AGO2 to confirm specificity of the antibody prior to use in immunohistochemistry (IHC) and immunofluorescence (IF). Further details are provided in Supplementary Table 1.

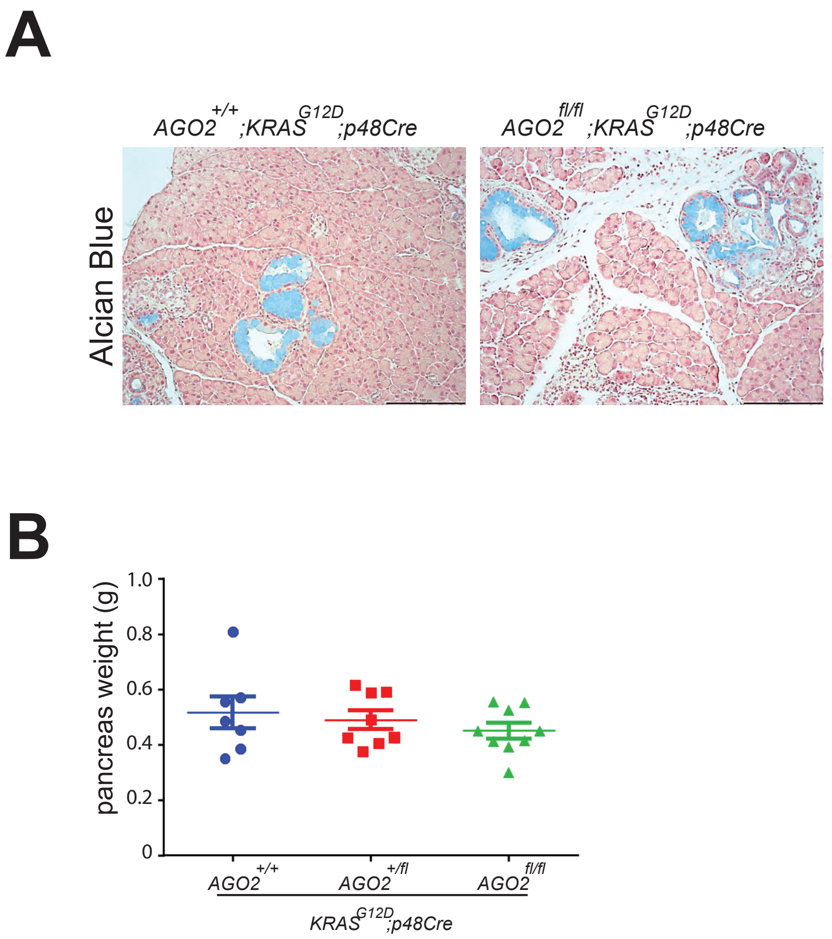


**Supplementary Figure 3: PanINs observed in *AGO2$^{+/+}$;KRAS$^{G12D}$;p48Cre* and *AGO2$^{fl/fl}$;KRAS$^{G12D}$;p48Cre* mice are similar.** (A) Alcian Blue (mucin) staining of PanINs of the indicated genotypes. (B) Scatter plot showing the weight of pancreata obtained from mice of three indicated genotypes at 12 weeks of age.

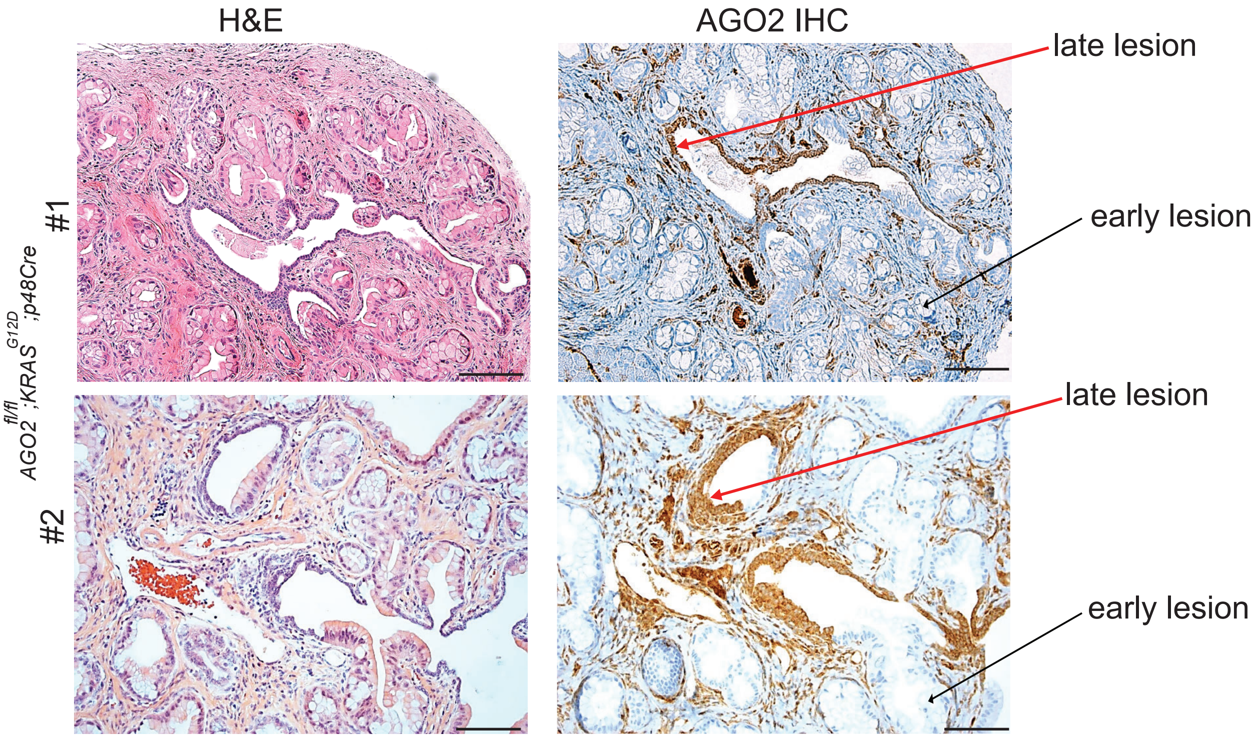


**Supplementary Figure 4: Leaky expression of AGO2.** Representative images of H&E and AGO2 IHC from two animals of the *AGO2*$^{fl/fl}$*;KRAS*$^{G12D}$*;p48Cre* cohort showing AGO2 expression in the late lesions. Scale bar represents 100 µm and 40 µm for upper and lower panels, respectively.

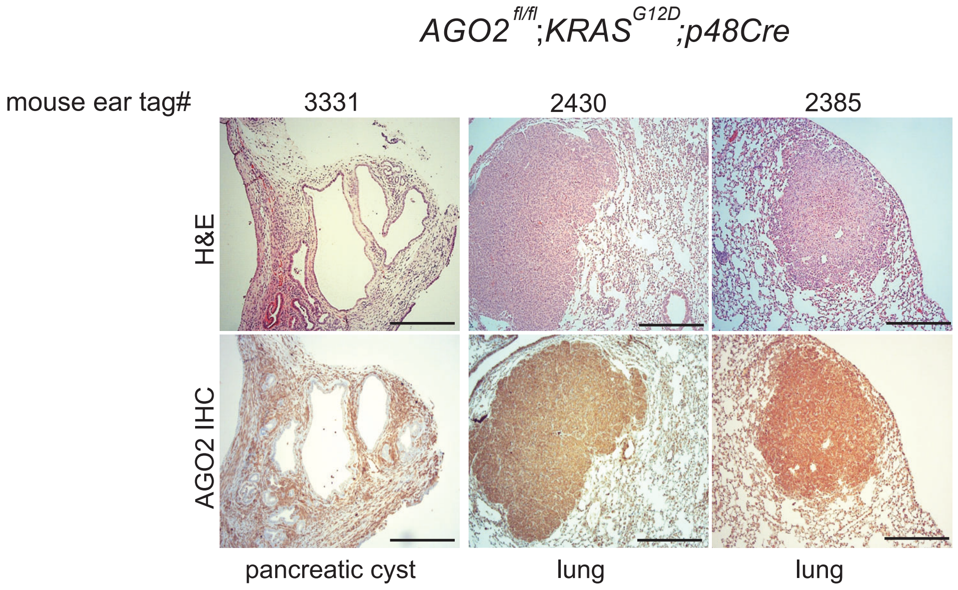


**Supplementary Figure 5: Abnormal pathologies in *AGO2* loss.** H&E and AGO2 IHC analysis of abnormal pancreas and lungs from the *AGO2*$^{fl/fl}$*;KRAS*$^{G12D}$*;p48Cre* mouse cohort (further detailed in Supplementary Table 2). Scale bar, 100 μm and 40 μm for low and high magnification, respectively.

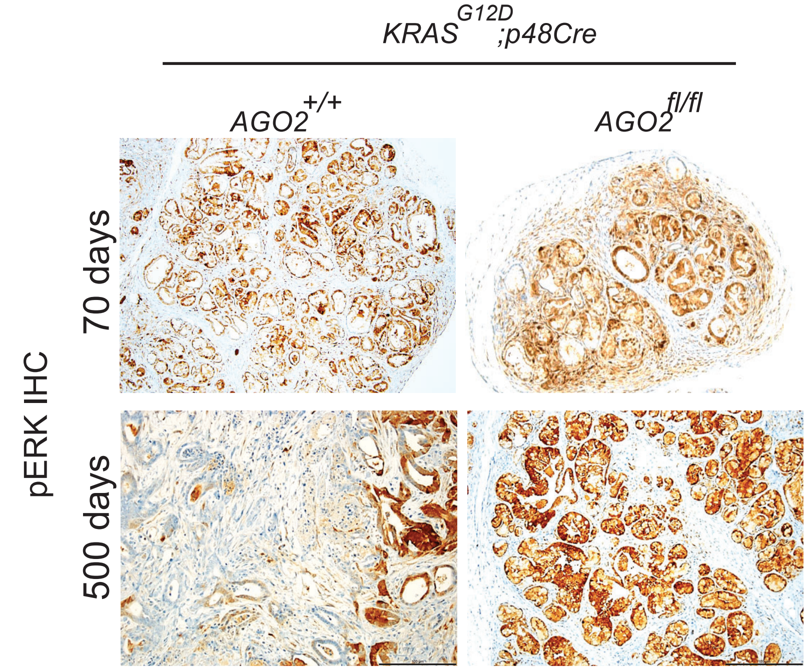


**Supplementary Figure 6: High phospho-ERK levels in PanINs of *AGO2$^{fl/fl}$;KRAS$^{G12D}$;p48Cre* mice.** Representative images of phospho-ERK IHC staining of pancreatic tissues from the indicated genotypes. Scale bar is 100 µm**.** Note the inconsistent staining pattern of phospho-ERK in PDAC tissue of *AGO2$^{+/+}$; KRAS$^{G12D}$;p48Cre* genotype at the 500-day time point.

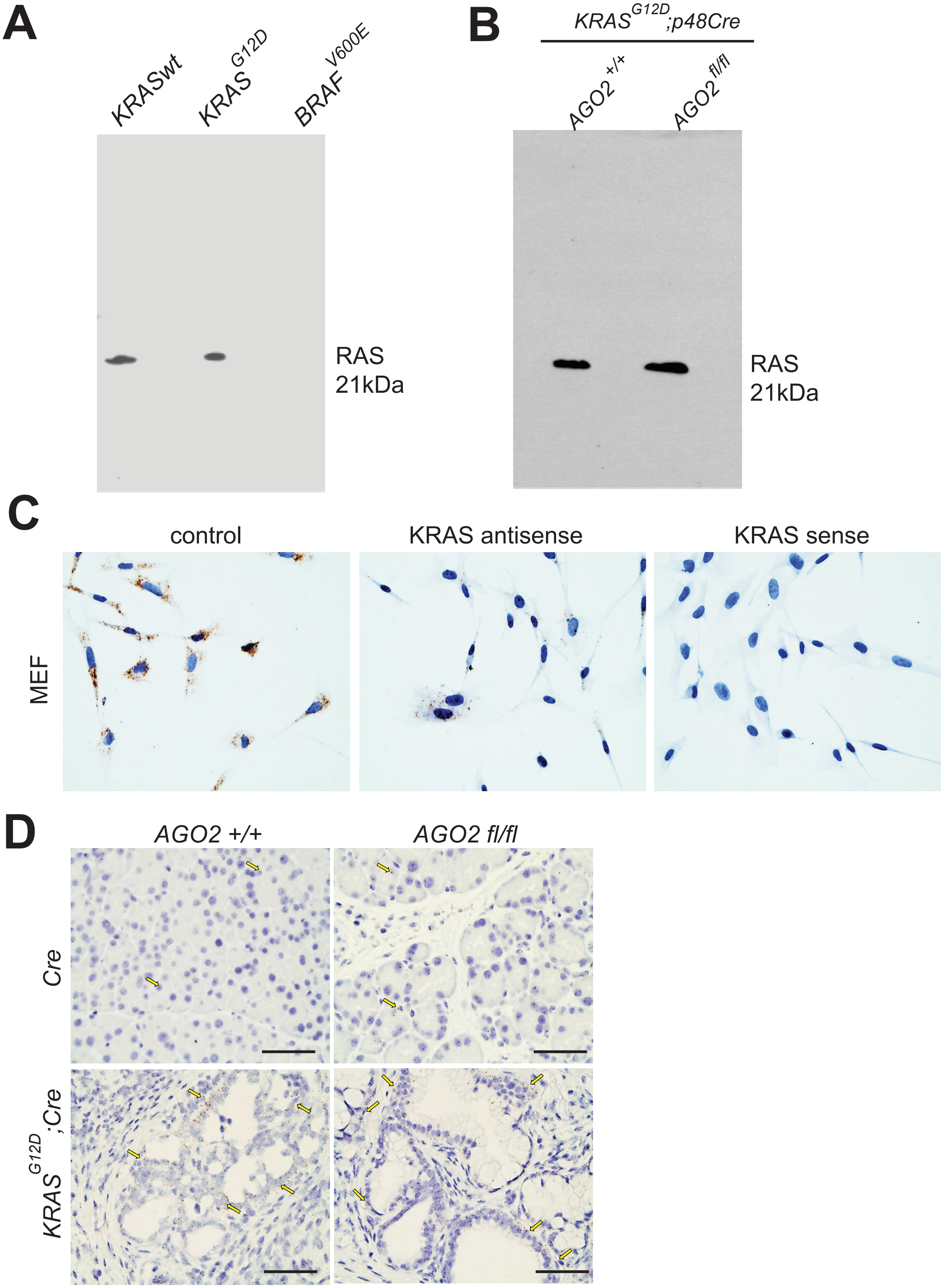


**Supplementary Figure 7: Specificity of RAS10 mab and *KRAS* transcript probes using RASless MEFs.** Specificity of the RAS10 antibody confirmed with immunoblot analysis of Rasless MEFs rescued using (A) various *KRAS* constructs and (B) mouse pancreas expressing oncogenic *KRAS*. Note that complete blots are shown to demonstrate the presence of a single band at 21 kDa. Further details are provided in Supplementary Table 1. (C) MEFs were assessed for *KRAS* RNA expression using RNA-ISH. *KRAS* antisense and sense probes were used as positive and negative controls, respectively. (D) Representative images of *KRAS* RNA-ISH on pancreatic tissues from normal and $AGO2^{fl/fl}$ mice (top panels) and oncogenic *KRAS*-expressing $AGO2^{+/+}$;$KRAS^{G12D}$;*Cre* and $AGO2^{fl/fl}$;$KRAS^{G12D}$;*Cre* mice (lower panels). Scale bar, 50 μm.

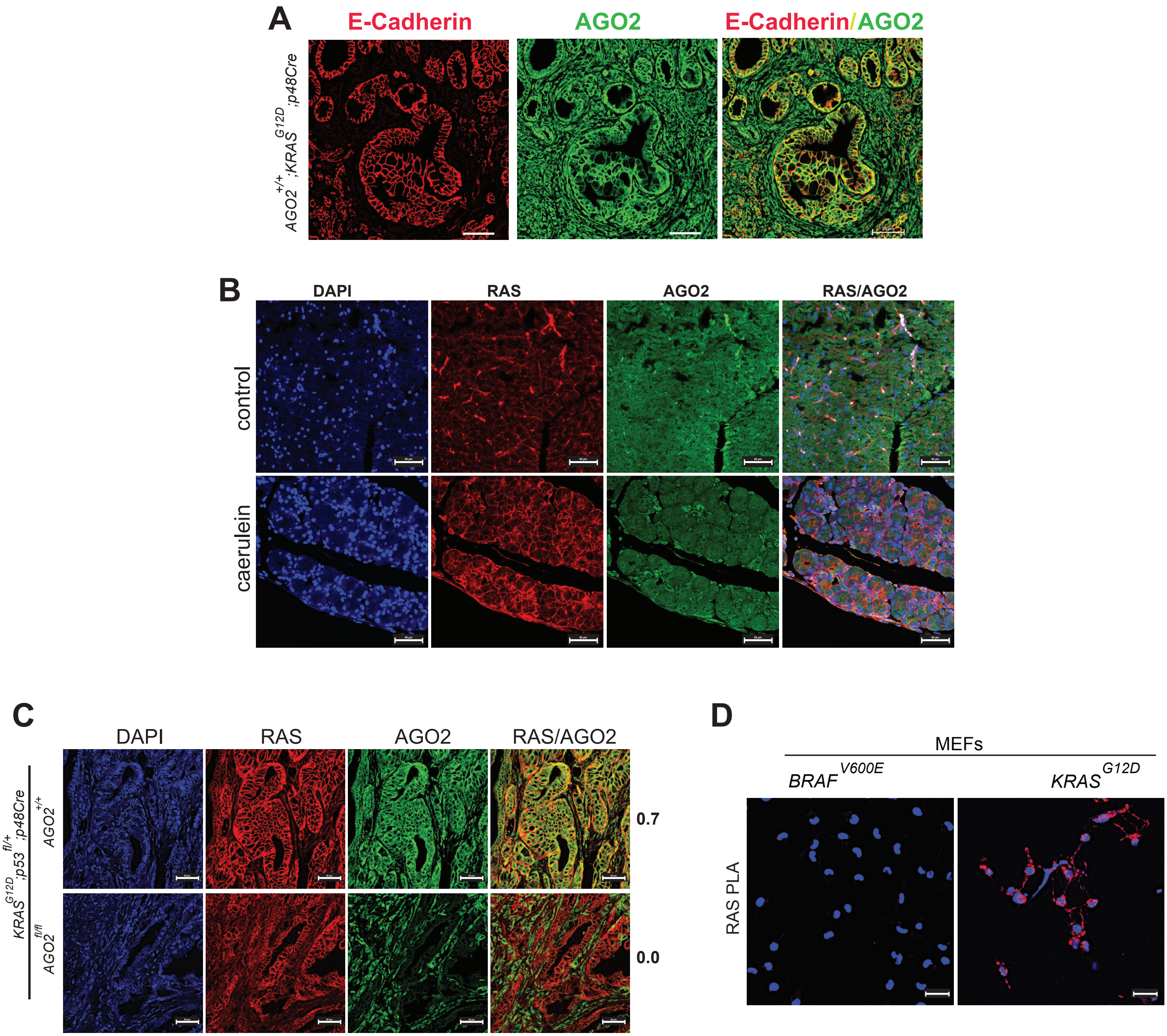


**Supplementary Figure 8: AGO2 membrane localization, RAS/AGO2 IF during pancreatitis or with p53 loss, and validation of RAS10 antibody for proximity ligation assay (PLA).** (A) Membrane localization of AGO2 demonstrated by AGO2/E-Cadherin co-localization signals in early PanIN lesions of *AGO2*$^{+/+}$;*KRAS*$^{G12D}$;*Cre* mice. E-Cadherin is used as a plasma membrane marker. Scale bar, 50 μm. (B) Representative images of RAS and AGO2 IF staining on pancreatic tissue from normal mice after treatment with caerulein to induce pancreatitis. (C) Representative images of IF analysis for RAS and AGO2 in PDAC lesions in pancreata of *AGO2*$^{+/+}$;*KRAS*$^{G12D}$;*p53*$^{fl/+}$;*Cre* and *AGO2*$^{fl/fl}$;*KRAS*$^{G12D}$;*p53*$^{fl/+}$;*Cre* mice. (D) Validation of RAS10 antibody for RAS PLA detection in RASless MEFs expressing either *KRAS*$^{G12D}$ or *BRAF*$^{V600E}$. Scale bar, 50 μm.

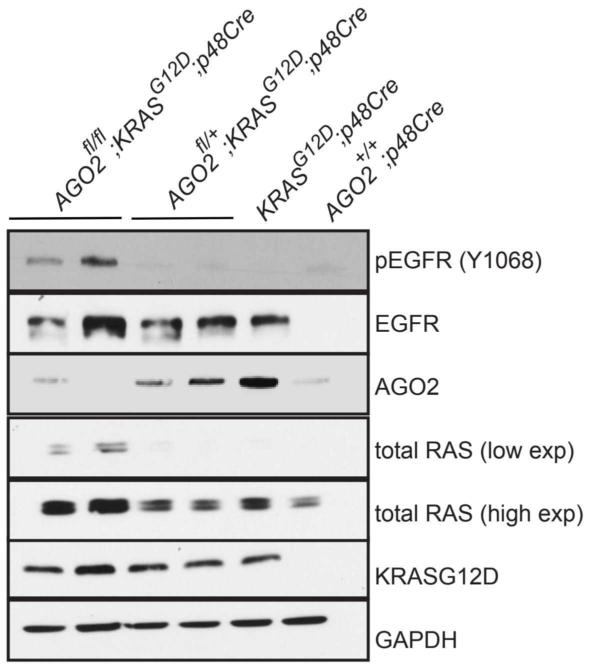


**Supplementary Figure 9: Immunoblot analysis of AGO2 and associated RAS signaling molecules from individual pancreata obtained from 12-week old mice of the indicated genotypes.**

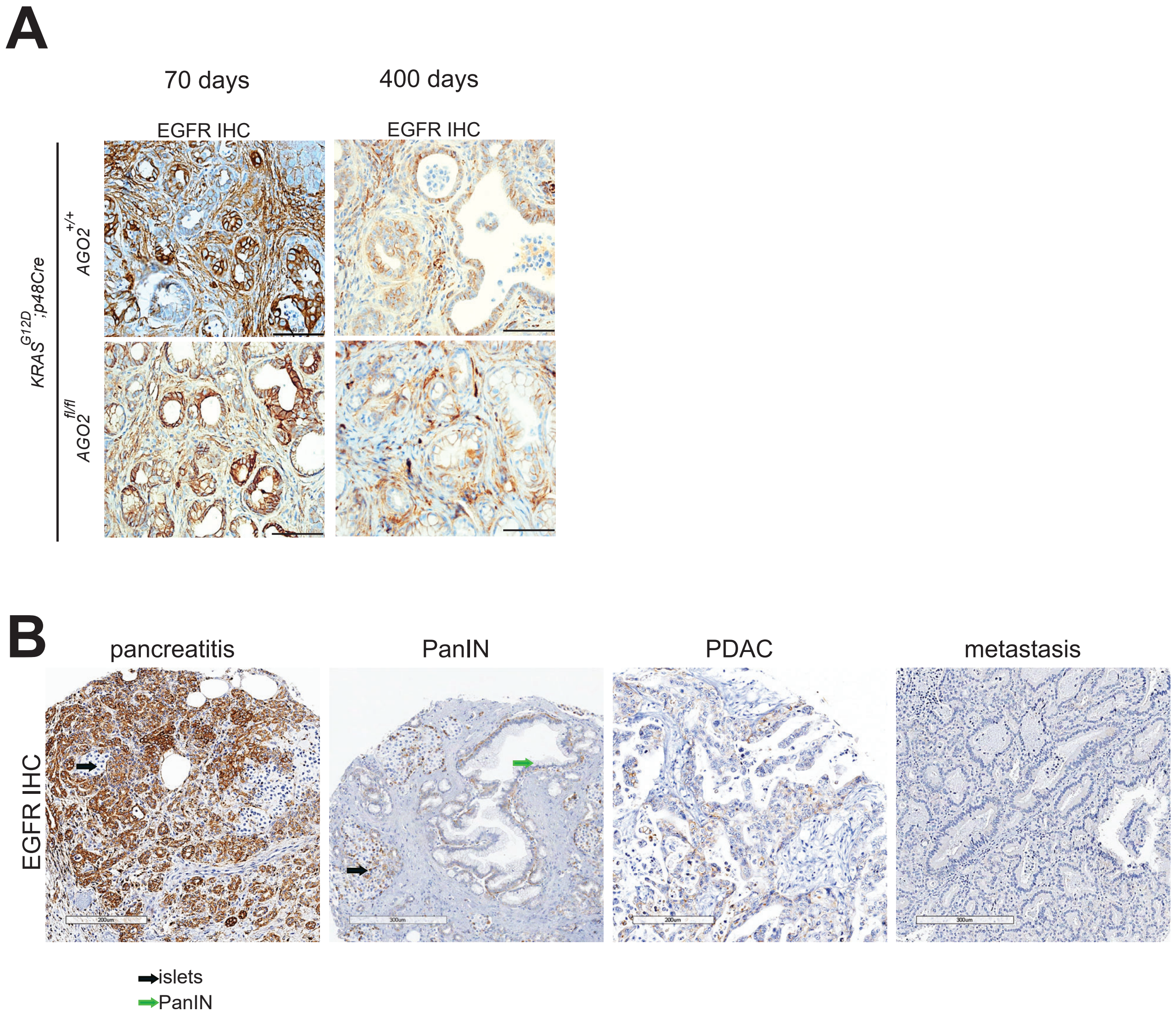


**Supplementary Figure 10: EGFR expression is reduced during pancreatic cancer progression.** (A) Representative images of EGFR IHC of pancreatic tissue from *AGO2*$^{+/+}$*;KRAS*$^{G12D}$*;p48Cre* and *AGO2*$^{fl/fl}$*;KRAS*$^{G12D}$*;p48Cre* mice at 70 or 400-day time points. Scale bar is 100 µm. (B) Representative images of EGFR IHC on a human pancreatic tissue microarray through disease progression. Scale bar is 300 µm.

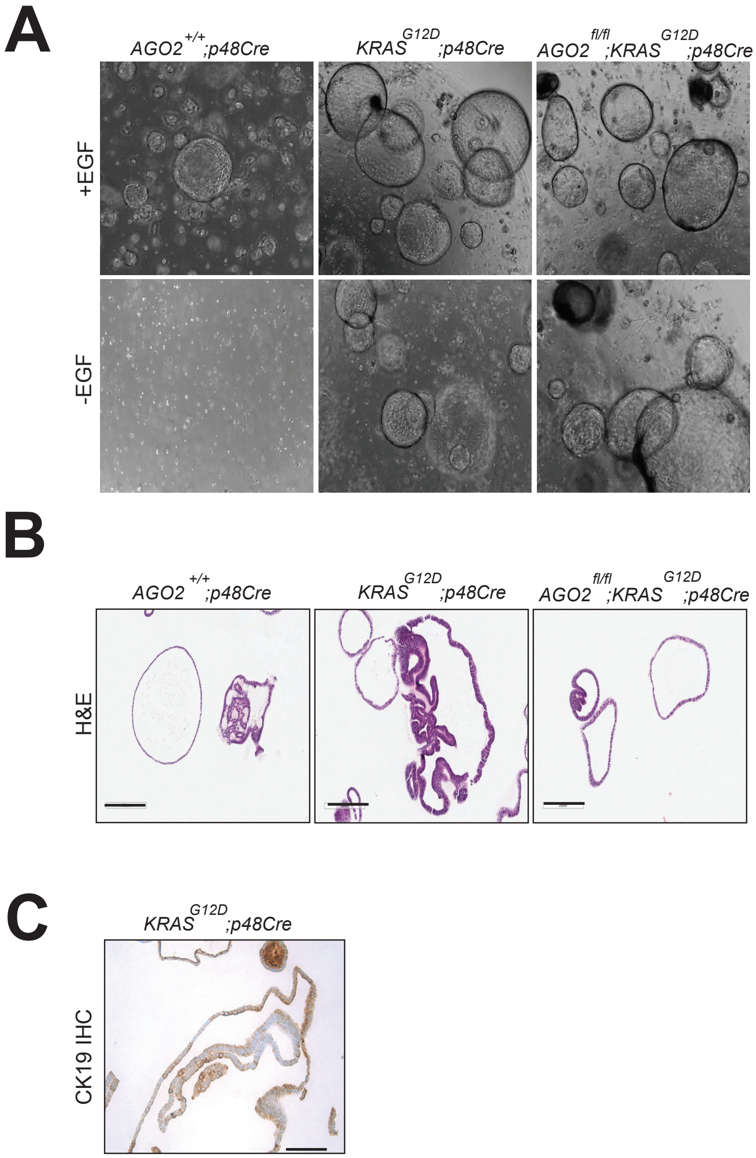


**Supplementary Figure 11: Characterization of pancreatic ductal organoids obtained from 12-week old *$AGO2^{+/+}$;$KRAS^{G12D}$;p48Cre* and *$AGO2^{fl/fl}$;$KRAS^{G12D}$;p48Cre* mice.** (A) Light microscopic images of mouse pancreatic ductal organoids obtained from the indicated genotypes and cultured in the presence or absence of EGF. (B) H&E staining of pancreatic ductal organoids. Scale bar, 200 μm. (C) CK19 (ductal marker) IHC staining of pancreatic ducts isolated from *$KRAS^{G12D}$;p48Cre* mouse. Scale bar, 100 μm.

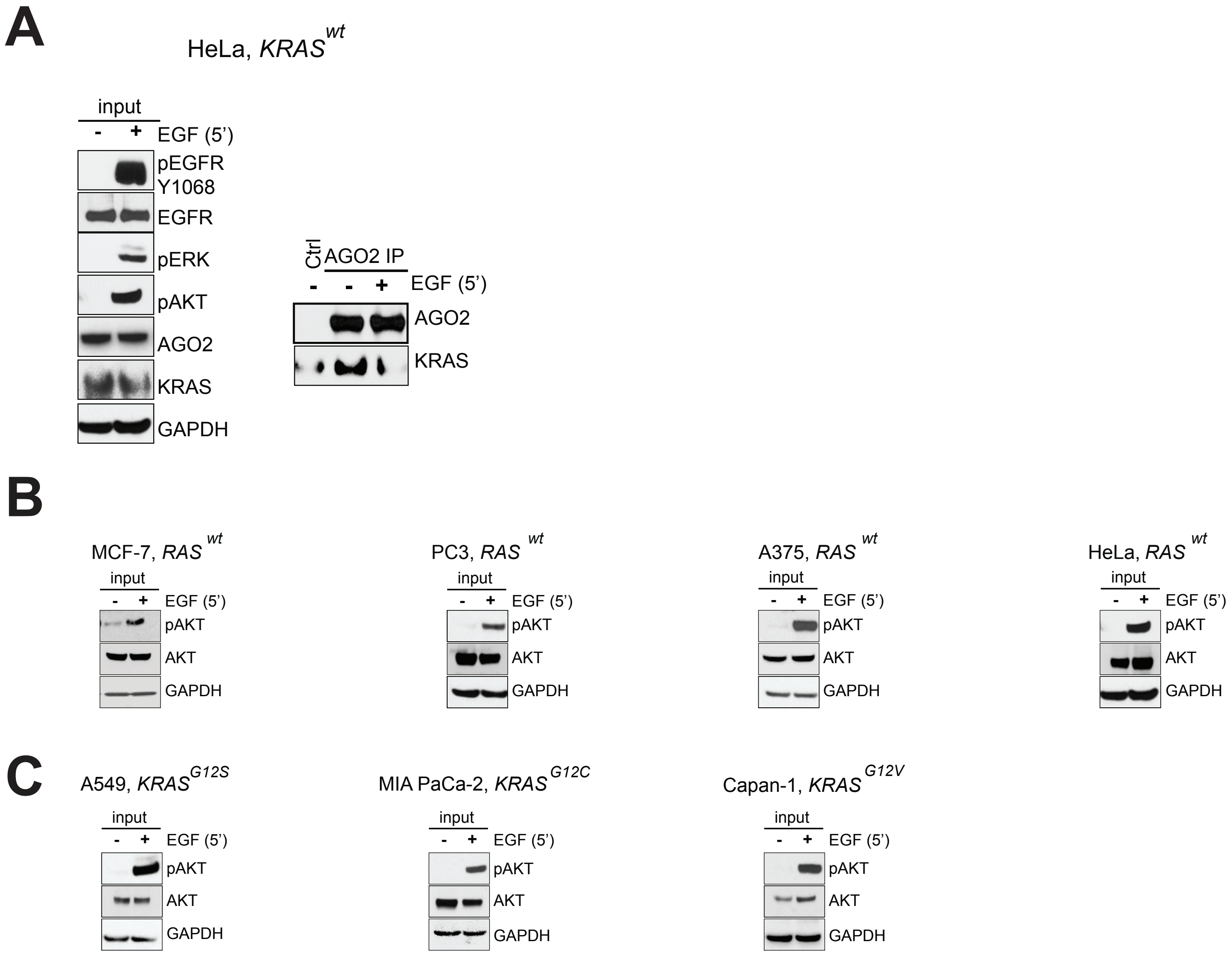


**Supplementary Figure 12: EGF stimulation disrupts the wild-type KRAS-AGO2 interaction.** (A) IP of endogenous AGO2 upon EGF stimulation (5'), in HeLa (*KRAS*$^{wt}$) cells followed by immunoblot analysis of KRAS. Input blots on the left show MAPK activation and levels of various proteins. (B) Input blots show Akt activation upon EGF stimulation in *KRAS*$^{wt}$ cells or (C) *KRAS*$^{mt}$ cells assessed in **Figure 7**.

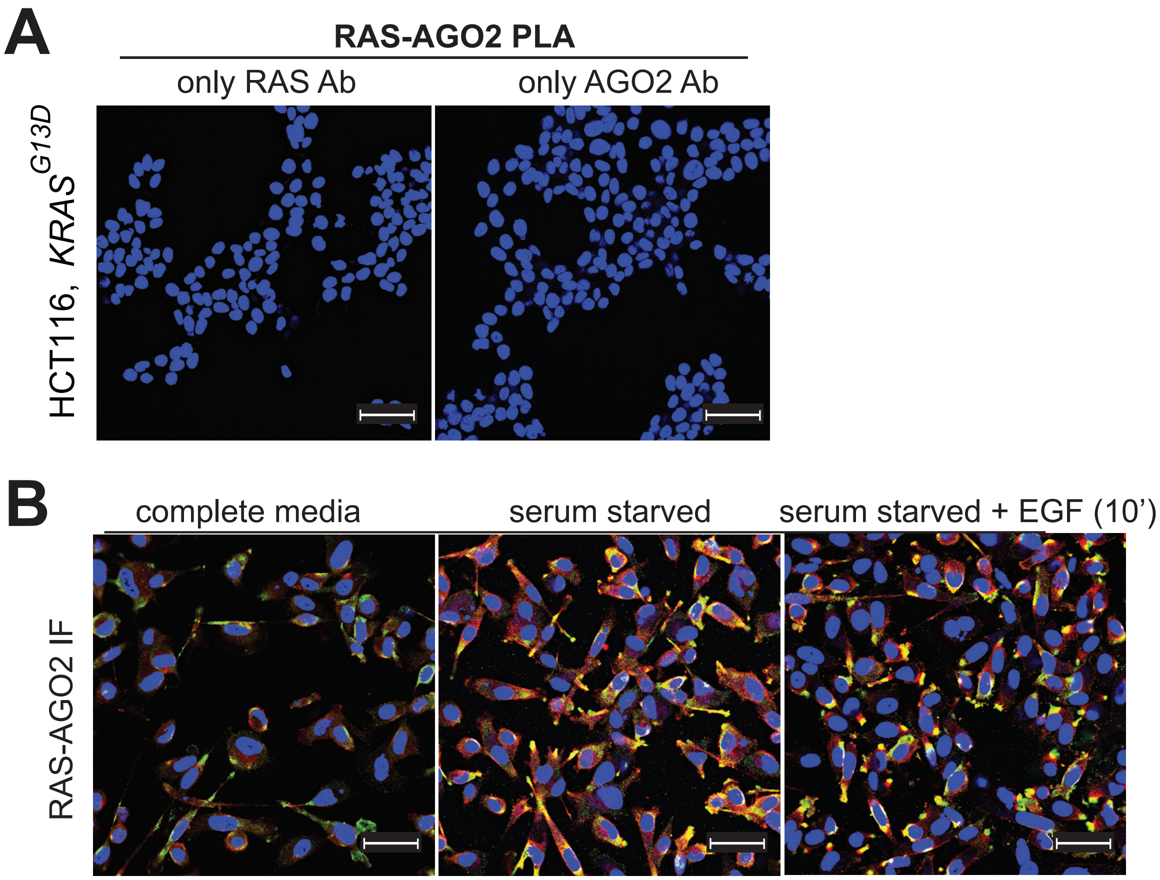


**Supplementary Figure 13: Validation of RAS-AGO2 PLA in cell line models.** (A) Control RAS-AGO2 PLA performed on HCT116 cells using either RAS or AGO2 antibody. (B) Representative images of RAS/AGO2 immunofluorescence in PC3 (*KRAS*$^{wt}$) cells grown under the indicated conditions. Confocal images were taken under identical settings across all cell culture conditions. Scale bar, 50 µm.
