## Supplemental Table 4 for "An Essential Role for *Argonaute 2* in EGFR-KRAS Signaling in Pancreatic Cancer Development"

**Supplementary Table 4: Cell lines used in the study**

| Number | Cell Line | Source | Tissue | Type | *KRAS* or *AGO2* Status |
|---|---|---|---|---|---|
| 1 | HEK293 | Human | Embryonic Kidney | Benign | *KRAS* WT |
| 2 | MCF-7 | Human | Breast | Cancer | *KRAS* WT |
| 3 | HeLa | Human | Cervix | Cancer | *KRAS* WT |
| 4 | A375 | Human | Skin Melanoma | Cancer | *KRAS* WT |
| 5 | PC3 | Human | Prostate | Cancer | *KRAS* WT |
| 6 | A549 | Human | Lung | Cancer | *KRAS* G12S |
| 7 | Capan-1 | Human | Pancreas | Cancer | *KRAS* G12V |
| 8 | MIA PaCa-2 | Human | Pancreas | Cancer | *KRAS* G12C |
| 9 | DLD-1 | Human | Colorectal | Cancer | *KRAS* G13D |
| 10 | MEF, parental | Mouse | Embryonic Fibroblast | Benign | *AGO2 WT* |
| 11 | MEF, *AGO2* -/- | Mouse | Embryonic Fibroblast | Benign | *AGO2* Knockout |
| 12 | MEF, *AGO2* -/- ; + *AGO2* | Mouse | Embryonic Fibroblast | Benign | MEF *AGO2-/-* + *AGO2* rescue |
| 13 | RASless MEF | Mouse | Embryonic Fibroblast | Benign | MEFs devoid of RAS |
