## Supplemental Table 1 for "An Essential Role for *Argonaute 2* in EGFR-KRAS Signaling in Pancreatic Cancer Development"

**Supplementary Table 1: Antibodies used for immunoprecipitation (IP), immunoblotting (IB), immunohistochemistry (IHC), and immunofluorescence (IF)**

| # | Antibody | Vendor | Catalog number | Application | Specificity | RAS antibody validation |
|---|---|---|---|---|---|---|
| 1 | Anti-Ras clone 10 (RAS10) | Millipore | 05-516 | IP, IB, IF | Human, mouse | This study & Waters et al., 2017 |
| 2 | K-Ras-2B antibody (C-19) | Santa Cruz | sc-521 | IP | Human | Waters et al., 2017 |
| 3 | K-Ras monoclonal antibody | Santa Cruz | sc-30 | IB | Human, mouse | Waters et al., 2017 |
| 4 | RAS (G12D mutant-specific) DH87 | Cell Signaling | 14429S | IB | Human | Waters et al., 2017 |
| 5 | AGO2, 11A9 | Sigma | SAB4200085 | IP, IB | Human | N/A |
| 6 | AGO2 EIF2C2 | Sino Biologicals | 11079-T36 | IB, IHC, IF | Human | N/A |
| 7 | Anti- EGFR ( phospho Y1092) | Abcam | ab40815 ( EP774Y) | IHC | Human, mouse | N/A |
| 8 | Phospho-EGF Receptor (Y1086) | Cell Signaling | 2220S | IB | Human, mouse | N/A |
| 9 | Phospho-p44/42 MAPK (Erk 1/2) | Cell Signaling | 4376 | IB | Human, mouse | N/A |
| 10 | ERK1(K-23) | Santa Cruz | sc-94 | IB | Human, mouse, rat | N/A |
| 11 | Total p44/42 MAPK (Erk 1/2) | Cell Signaling | 9102 | IB | Human, mouse | N/A |
| 12 | EGFR antibody | Abcam | ab52894 ( EP38Y) | IB,IHC | Human, mouse, rat | N/A |
| 13 | EGFR(1005) | Santa Cruz | sc-03 | IB | Human, mouse, rat | N/A |
| 14 | Anti-FLAG antibody | Sigma | F7425-0.2MG | IP, IB | Human, mouse | N/A |
| 15 | Phospho-Akt S473 | Cell Signaling | 4060S | IB | Human,mouse | N/A |
| 16 | E-cadherin (36) | Ventana Roche | 790-4497 | IF | Mouse | N/A |
| 17 | AKT (pan) - C67E7 | Cell Signaling | 4691 | IB | Human,mouse | N/A |
| 18 | Anti-Cytokeratin 19 antibody | Abcam | ab133496 | IHC | Mouse | N/A |
| 19 | GAPDH-HRP | Cell Signaling | 3683 | IB | Human, mouse | N/A |
| 20 | Normal mouse IgG | Santa Cruz | sc-2025 | Isotype control IP | Mouse | N/A |
| 21 | Normal rat IgG | Abcam | ab18450 | Isotype control IP | Rat | N/A |
| 22 | Normal rabbit IgG | Millipore | 12-370 | Isotype control IP | Rabbit | N/A |
