## Supplemental Table 2 for "An Essential Role for *Argonaute 2* in EGFR-KRAS Signaling in Pancreatic Cancer Development"

**Supplementary Table 2: Cases in *AGO2$^{fl/fl}$;KRAS$^{G12D}$;p48Cre* cohort with abnormal pathologies or death prior to 500-day time point**

| | | | |
|---|---|---|---|
| Ear tag# | 3331 | 2430 | 2385 |
| Age (days) | 368 | 500 | 500 |
| Liver | | | |
| Ascites | | | |
| Lungs | | | |
| Diaphragm | | | |
| Hemorrhage | | | |
| Lymph nodes | | | |
| Kidneys | | | |
| Jaundice | | | |
| Enlarged spleen | | | |
| **Pathological Evaluation** | | | |
| PDAC | No anaplastic PDAC or PDAC | No PDAC | No PDAC |
| Pancreas | Low grade PanINs and cyst | Only low grade PanIN | Only low grade PanIN |
| AGO2 expression in pancreas | No AGO2 expression in the cyst or PanINs | No AGO2 expression in early PanINs | No AGO2 expression in early PanINs |
| Pancreatic cyst | Yes | No cyst | No cyst |
| Further comment/ other pathology | Resembles mucinous cystic neoplasm | Likely benign lesion in lung, unknown origin | Likely benign lesion in lung, unknown origin |
| AGO2 expression in mets? | N/A | AGO2 expressed in benign lung lesion | AGO2 expressed in benign lung lesion |
| Cause of death | Unknown | Not dead before 500 days | Not dead before 500 days |
