## Supplemental Table 3 for "An Essential Role for *Argonaute 2* in EGFR-KRAS Signaling in Pancreatic Cancer Development"

**Supplementary Table 3: Primers used in the study**

| Transcript | Primer Sequence |
|---|---|
| m*HRAS*_Fwd | *5' GCTTCCTCTGTGTATTTGCCA 3'* |
| m*HRAS*_Rev | *5' CTTTCACCCGCTTGATCTGC 3'* |
| m*KRAS*_Fwd | *5' GTTAGCTCCAGTGCCCCAAT 3'* |
| m*KRAS*_Rev | *5' ATTCCCTAGGTCAGCGCAAC 3'* |
| m*NRAS*_Fwd | *5' ACTGGCCAAGAGTTACGGAA 3'* |
| m*NRAS*_Rev | *5' TGGCGTATCTCCCTTACCAG 3'* |
| m*AGO2*_Fwd | *5' GATCGCCAAGAGGAGATCAG 3'* |
| m*AGO2*_Rev | *5' GCCTCCCAGTTTGACATTGA 3'* |
| m*GAPDH*_Fwd | *5' AAGGTCATCCCAGAGCTGAA 3'* |
| m*GAPDH*_Rev | *5' CTGCTTCACCACCTTCTTGA 3'* |
| hAGO2 ORF Fwd | *5' ATCAACGTCAAGCTGGGAGG 3'* |
| hAGO2 ORF Rev | *5' GTGACGTCTGCTCCCAGAAA 3'* |
